# The KASH protein UNC-83 differentially regulates kinesin-1 activity to control developmental stage-specific nuclear migration

**DOI:** 10.1101/2025.03.06.641899

**Authors:** Selin Gümüşderelioğlu, Natalie Sahabandu, Daniel Elnatan, Ellen F. Gregory, Kyoko Chiba, Shinsuke Niwa, G.W. Gant Luxton, Richard J. McKenney, Daniel A. Starr

## Abstract

Nuclear migration plays a fundamental role in development, requiring precise spatiotemporal control of bidirectional movement through dynein and kinesin motors. Here, we uncover a mechanism for developmental regulation of nuclear migration directionality. The nuclear envelope KASH protein UNC-83 in *Caenorhabditis elegans* exists in multiple isoforms that differentially control motor activity. The shorter UNC-83c isoform promotes kinesin-1-dependent nuclear movement in embryonic hyp7 precursors, while longer UNC-83a/b isoforms facilitate dynein-mediated nuclear migration in larval P cells. We demonstrate that UNC-83a’s N-terminal domain functions as a kinesin-1 inhibitory module by directly binding kinesin heavy chain (UNC-116). This isoform-specific inhibition, combined with differential affinity for kinesin light chain (KLC-2), establishes a molecular switch for directional control. Together, these interdisciplinary studies reveal how alternative isoforms of cargo adaptors can generate developmental stage-specific regulation of motor activity during development.

## INTRODUCTION

Nuclear positioning orchestrates essential cellular and developmental processes including pronuclear migration, muscle and neuronal differentiation, and cell migration^1,2,3,4^. Defects in nuclear positioning underlie severe congenital disorders such as lissencephaly^5,6^ and centronuclear myopathies^7^. As the largest intracellular cargo, nuclei require precise spatial and temporal control of their movement along microtubules^4^. Nuclei move towards different directions along microtubules at different times in development. For example, the minus-end directed motor cytoplasmic dynein (dynein from here on) drives pronuclear migration toward centrosomes during fertilization ^8,9,10,11,12^. Alternatively, the plus-end directed motor kinesin powers nuclear movement in developing neuroepithelia and sensory cells^1,13^. However, the molecular mechanisms controlling bidirectional nuclear movements remain poorly understood. We hypothesize that nuclear envelope cargo adaptors regulate bidirectional nuclear movements by directly influencing the activity of kinesin and dynein motors.

The conserved **<u>li</u>**nker of the **<u>n</u>**ucleoskeleton and **<u>c</u>**ytoskeleton (LINC) complex bridges the nuclear envelope to enable many nuclear movements^14,15,16^. LINC complexes are formed by the interaction between **<u>S</u>**ad1p/**<u>UN</u>**C-84 (SUN) proteins in the inner nuclear membrane and **<u>K</u>**larsicht/**<u>A</u>**NC-1/**<u>S</u>**yne **<u>h</u>**omology (KASH) proteins in the outer nuclear membrane^17,18^. SUN proteins interact with lamins in the nucleoplasm^19,20^, cross the inner nuclear membrane, span the lumen of the nuclear envelope, and recruit KASH proteins to the outer nuclear membrane^16,21^. KASH proteins, including the mammalian nuclear envelope spectrin repeat proteins (nesprins), extend into the cytoplasm to interact with a variety of cytoskeletal and motor proteins ^22,23,24^. Here we focus on the role of KASH proteins as cargo adaptors to recruit kinesin-1 and dynein motors to nuclei.

Motor protein recruitment by KASH proteins involves complex regulatory mechanisms. Kinesins are primarily plus-end directed motors while cytoplasmic dynein is the major minus-end directed motor in animal cells^25,26^. Kinesin-1 is a heterotetramer composed of two heavy chains (KHCs) with their motor domains and two kinesin light chains (KLCs)^25,27,28^. In the absence of cargo, kinesin-1 is inactive as KHCs and KLCs are folded in a confirmation that inhibits microtubule binding and motility^29,30^. Binding of cargo adaptor molecules to the KLC releases this autoinhibition and activates kinesin-1 motility^31,32,33,34,35^. Mammalian nesprin-2 and nesprin-4, and *C. elegans* UNC-83 activate kinesin-1 through their W-acidic (LEWD/EWD) motifs that interact with the tetratricopeptide repeat (TPR) domain of KLCs^13,36,37,38^. Similarly, dynein exists in an autoinhibited state until activated by co-factors and cargo adaptors^26,39,40^. At the nuclear envelope, mammalian KASH5, and *C. elegans* ZYG-12 are meiosis-specific activators of dynein^41,42,43,44^. Nesprin-1 and –2 also bind dynein^45^. Despite our understanding of these individual interactions, the mechanisms controlling tissue-specific activation of different motor proteins by KASH proteins remains unclear.

We studied two distinct nuclear migration events in *C. elegans* development to dissect how KASH proteins coordinate microtubule motor protein regulation. First, embryonic hypodermal hyp7 precursor nuclei migrate across the dorsal midline to the opposite lateral side of the embryo, after which the cells fuse to form the hyp7 syncytium^46^ (Fig. 1A). Failed migration results in nuclei mislocalized to the dorsal cord^15,47^. In contrast, larval hypodermal P-cell nuclei migrate ventrally through constricted spaces between body wall muscles and the cuticle. After migration, P cells undergo divisions to ultimately form vulval cells and GABA neurons^48,49^ (Fig. 1B). Failures in P-cell nuclear migration result in P-cell death and missing lineages that lead to egg-laying and locomotion defects^49,50^.

**Figure 1:**
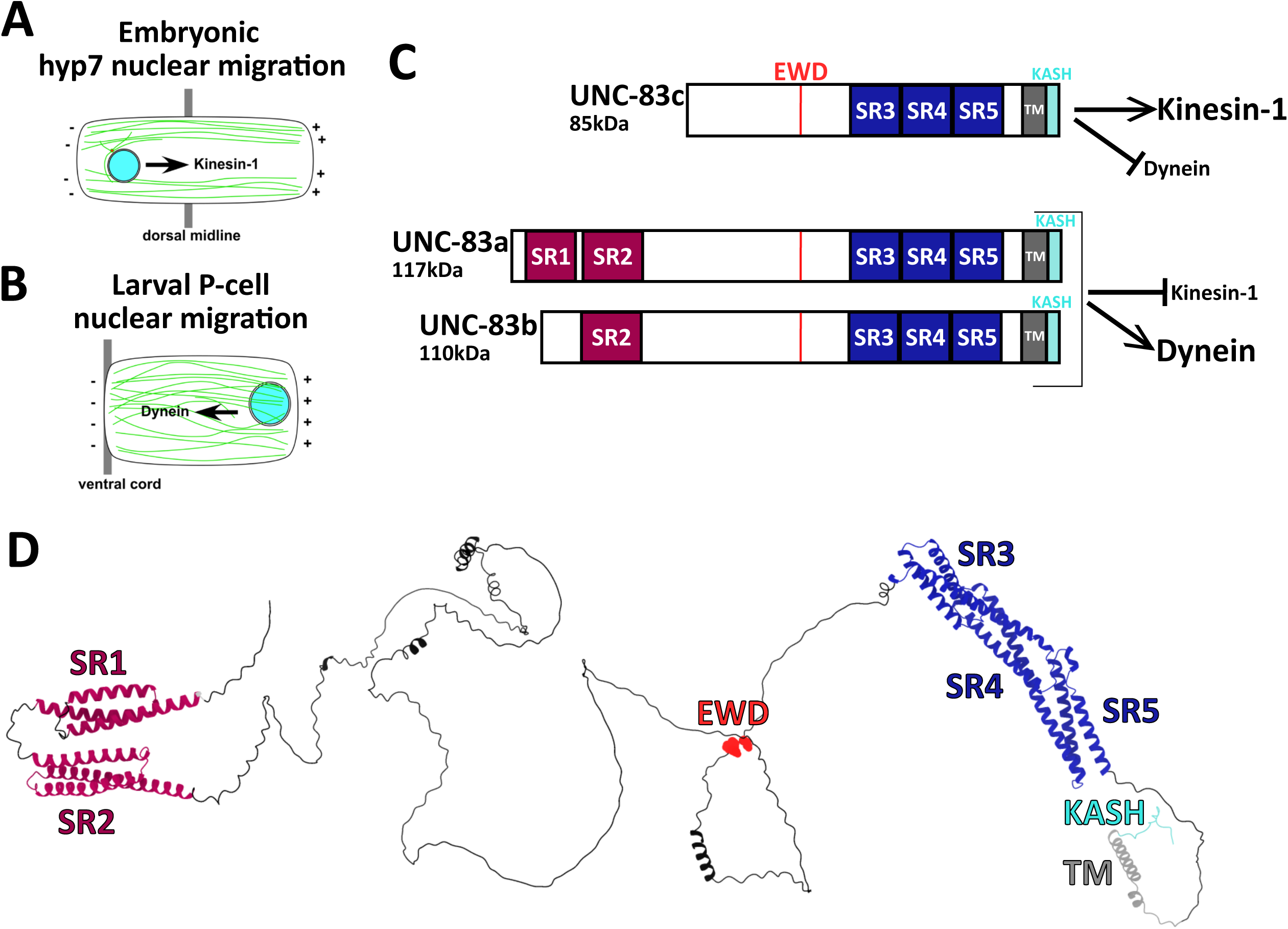
Different isoforms of UNC-83 mediate nuclear migration throughout development. A) Schematic showing the kinesin-1-mediated migration of embryonic hypodermal hyp7 nuclei (cyan) across the dorsal midline (gray) towards the plus-ends of microtubules (green). B) Schematic illustrating the dynein-driven migration of larval P-cell nuclei (cyan) towards the ventral cord (gray), where the minus-ends of microtubules are positioned. C) Model of the hypothesis tested in this study. The short isoform UNC-83c activates kinesin-1 and/or inhibit dynein, making kinesin-1 the main motor driving embryonic hyp7 nuclear migration. The longer isoforms UNC-83a/b activate dynein and/or inhibit kinesin-1, making dynein the main driver of larval P-cell nuclear migration. D) AlphaFold3 predicted structure of the long isoform, UNC-83a. C-D) Spectrin-like repeats (SRs) SR1 and SR2 (unique to UNC-83a/b) are in magenta; SR3, SR4 and SR5 (common to all isoforms) are in dark blue. The transmembrane domain (TM) is shown in dark gray. The KASH peptide is shown in cyan. The EWD motif is highlighted in red (see Figure S1 for confidence and error scores of the predicted structure).

Both nuclear migration events require the LINC complex proteins UNC-84 (SUN) and UNC-83 (KASH) ^51,52,53,54^. UNC-83 functions as a cargo adaptor that directly interacts with both kinesin-1 and dynein at the nuclear envelope^54,55,56^. However, cellular and genetic analyses reveal strict tissue-specific motor requirements. Embryonic hyp7 precursor nuclei migrate toward the plus ends of microtubules (Fig. 1A), while P-cell nuclei migrate toward minus ends of microtubules (Fig. 1B)^15,49^. Mutations in kinesin-1 heavy chain (*unc-116*) or light chain (*klc-2*) severely disrupt hyp7 nuclear migration, while dynein (*dhc-1*) plays only a minor role to move nuclei backward to avoid obstacles ^55,56^. In contrast, dynein, but not kinesin-1, is necessary for P-cell nuclear migration^49,57^. This precise tissue specificity in motor preference suggests that UNC-83 can differentially recruit and/or activate kinesin-1 for plus-end-directed hyp7 nuclear migration and dynein for minus-end-directed P-cell nuclear migration.

We hypothesized that the differential expression of *unc-83* isoforms controls tissue-specific nuclear migration directionality. There are three known *unc-83* isoforms; *unc-83a*, *unc-83b*, and *unc-83c*, encoding proteins with 1041, 974 and 741 amino acid residues, respectively. The three isoforms have 741 residues in their C-termini in common, but differ in their N-termini^52^. Previous genetic analyses revealed that mutations affecting all three isoforms disrupt both migration events, while those specific to long isoforms (*unc-83a/b*) only impair P-cell nuclear migration. Therefore, the short isoform (*unc-83c*) suffices for hyp7 nuclear migration^52^. In this model, the long UNC-83a/b isoforms favor minus-end-directed nuclear migration by inhibiting kinesin-1 and/or activating dynein while the short UNC-83c isoform favors plus-end-directed movements by activating kinesin-1 and/or inhibiting dynein (Fig. 1C). Here, we test our model with *in vivo* and *in vitro* approaches. Together, our findings reveal that the N-terminus of long UNC-83 isoforms specifically inhibits kinesin-1, enabling precise spatial and temporal control of nuclear positioning during development.

## RESULTS

### AlphaFold predicts a modular spectrin-repeat architecture for UNC-83

The molecular architecture of UNC-83’s cytoplasmic domain remains unexplored. Given that UNC-83 is predicted to be highly α-helical and that other KASH proteins contain characteristic ∼100 residue long spectrin repeats composed of triple α-helical bundles^56,58^, we hypothesized that UNC-83 might harbor previously unidentified spectrin-like domains. AlphaFold3-mediated structural modeling^59^ revealed five spectrin-like repeats interspersed by large stretches of predicted disordered regions (Fig. 1D). Two spectrin-like repeats were located within the N-terminal domain specific to the longer UNC-83a/b isoforms while the other three were located within the C-terminal region shared by all isoforms (Fig. 1C-D). The KLC-binding EWD motif was in an unstructured region (Fig.1D), similar to nepsrin-2^60^. These structural predictions show high confidence scores (Fig. S1A) and low predicted error rates (Fig. S1B), suggesting that UNC-83, like other KASH proteins, evolved from an ancestral spectrin-like protein.

### The N-terminal domain of the long UNC-83a isoform inhibits KLC-2 binding

Previous studies established that the short UNC-83c isoform directly binds KLC-2 via yeast two hybrid and *in vitro* pull-down assays^55^, and that both proteins are required for plus-end directed nuclear migration in embryonic hyp7 precursors^15^. The conserved EWD motif of UNC-83 is predicted to interact with the TPR repeats of KLCs as modeled by AlphaFold3 (Fig. 2A) and as demonstrated for other kinesin-1 cargo adaptors^36^. The importance of this motif is supported by the *unc-83(yc74)* allele, where mutation of EWD to GSA residues abolishes hyp7 nuclear migration^13^. While this EWD motif is present in all three UNC-83 isoforms and is necessary for kinesin-1-mediated hyp7 nuclear migration, we hypothesized that the 301-residue N-terminal portion unique to UNC-83a/b isoforms inhibits binding to KLC-2.

**Figure 2:**
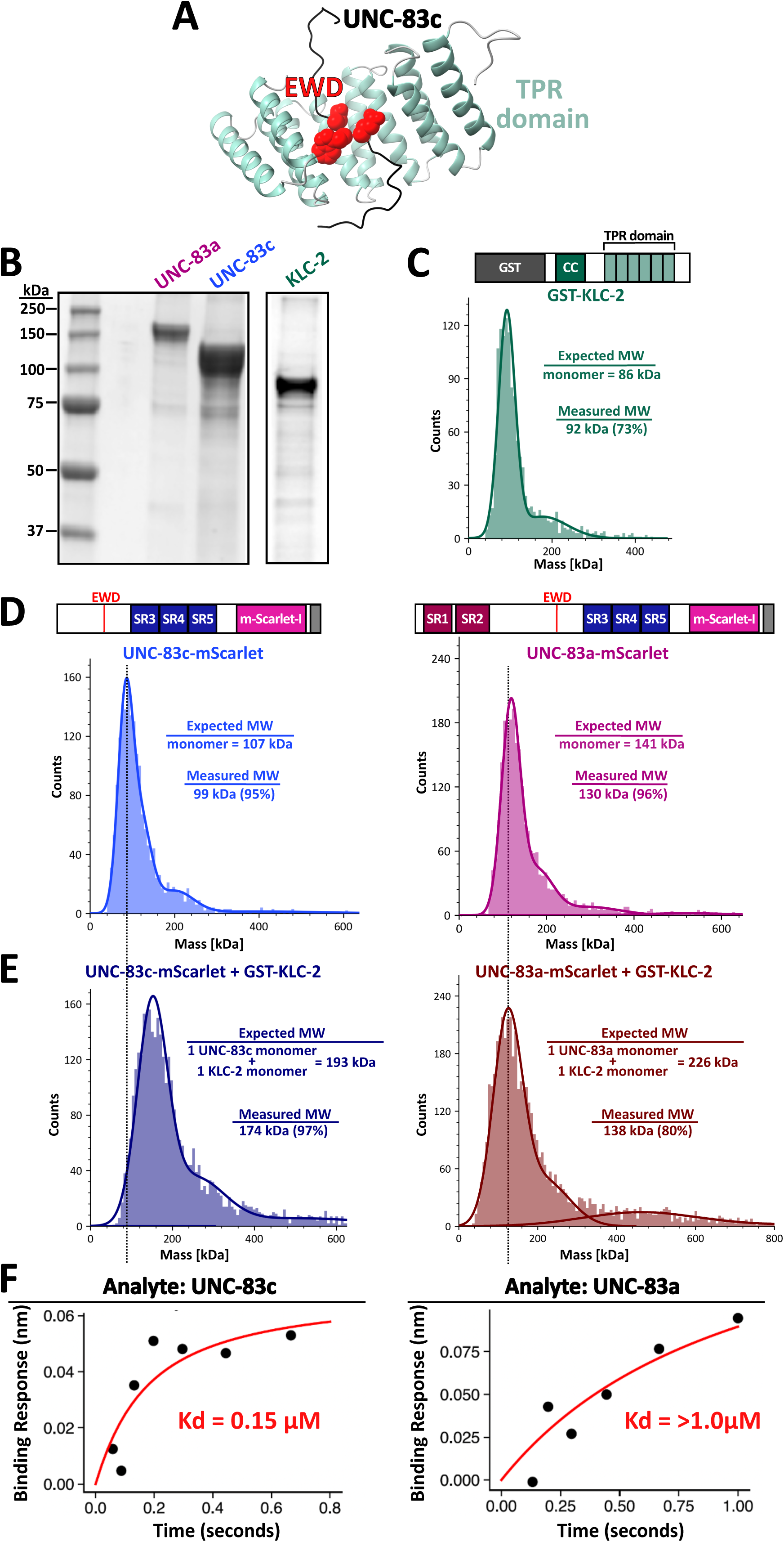
The N-terminal domain of the long UNC-83a isoform inhibits KLC-2 binding. A) AlphaFold3-predicted interaction of UNC-83c’s EWD motif (red) with the TPR domain (green) of the kinesin light chain, KLC-2. B) Coomassie-stained protein gels of purified UNC-83a, UNC-83c and KLC-2 recombinant proteins. C) Schematic of the GST-KLC-2 construct (GST tag in gray, coiled-coil domain in dark green, and TPR domain in light green) and a Gaussian fit of its measured molecular weight in solution (30 nM; monomer = 92 + 19.8 kDa, 72% peak population; dimer = 177 + 59 kDa, 21% peak) as determined by mass photometry. D) Schematics and molecular weight distributions of UNC-83c-mScarlet (30 nM; monomer = 99 + 47 kDa, 95% peak) and UNC-83a-mScarlet (30 nM; monomer = 130 + 62 kDa, 96% peak). SR1 and SR2 (unique to UNC-83a/b) are in magenta; SR3, SR4 and SR5 (common to all isoforms) are in dark blue; mScarlet-I tag in fuchsia, and Strep-II tag in dark gray. E) Molecular weight distributions of KLC-2 preincubated with UNC-83c (30 nM; one UNC-83c monomer + one KLC-2 monomer = 174 + 134 kDa, 97% peak) and UNC-83a (30 nM; one UNC-83a monomer = 138 + 62 kDa, 80% peak). F) Binding curves of UNC-83a and UNC-83c to KLC-2, showing an affinity of 0.15 µM for UNC-83c and >1.0 µM for UNC-83a. The bait, GST-KLC-2, was diluted to 1 µM and immobilized onto the biosensors. Both UNC-83a and UNC-83c were tested at concentrations of 1.0, 0.667, 0.444, 0.296, 0.197, 0.132, 0.088 μM.

To test our hypothesis and quantitatively assess UNC-83c isoform binding to KLC-2, we expressed and purified KLC-2 and the cytoplasmic domains of UNC-83a and UNC-83c (excluding their transmembrane domains and KASH peptides for solubility) from *E. coli* (Fig. 2B). Mass photometry analysis^61^ confirmed that all three purified proteins were predominantly monomeric, with measured masses within 10% of their predicted molecular weights (Fig. 2C-D). Direct binding assays revealed that UNC-83c and KLC-2 formed a stable complex (one UNC-83c monomer binding to one KLC-2 monomer) with an average mass of 174 kDa (Fig. 2E). In contrast, UNC-83a did not form a robust complex with KLC-2 under identical conditions (Fig. 2E). While a minor peak at ∼489 kDa was observed, this represented less than 20% of the detected particles and likely reflects transient or non-specific associations rather than a stable complex formation. These results suggest the N-terminal domain of UNC-83a modulates the ability of the EWD motif to interact with KLC-2.

To further quantify these differences in binding affinity, we used bio-layer interferometry for real-time interaction analysis. UNC-83c exhibited robust binding to KLC-2 with a dissociation constant (K_d_) of 0.15 µM. However, UNC-83a binding to KLC-2 showed non-saturating kinetics indicating a K_d_ at least 10-fold higher (K_d_ > 1.0) than that for UNC-83c (Fig. 2F). We conclude that the N-terminal domain specific to UNC-83a disrupts the interaction between its EWD motif and the TPR domain of KLC-2.

### Ectopic expression of *unc-83a* disrupts kinesin-1-dependent nuclear migration

Our biochemical data showed that the N-terminal domain of UNC-83a disrupts KLC-2 binding *in vitro*. To test whether this inhibitory effect extends to nuclear migration *in vivo*, we engineered *unc-83(yc108[UNC-83a])* using CRISPR/Cas9 to express the long UNC-83a isoform under control of the endogenous *unc-83c* promoter. This was achieved by fusing the 301-amino acid UNC-83a-specific N-terminal domain to the 5’ end of *unc-83c* (Fig. 3A), effectively replacing the short isoform with UNC-83a in hyp7 precursor cells. We confirmed the proper localization of UNC-83a to the nuclear envelope of hyp7 precursors during the nuclear migration period in *unc-83(yc108[UNC-83a])* by immunofluorescence (Fig. 3B). To asses nuclear migration defects, we quantified hyp7 nuclei abnormally localized in the dorsal cord of larvae^62^. Wild-type animals had almost no mislocalized hyp7 nuclei. In contrast, both *unc-83(e1408)* null and *unc-83(yc108[UNC-83a])* animals had severe defects with 12.8+ 0.45 and 12.9 + 0.32 (mean + 95% confidence interval (CI)) mislocalized nuclei, respectively (Fig. 3C-D). Importantly, *unc-83(yc108[UNC-83a])* animals had normal P-cell nuclear migrations, with only 0.45 + 0.16 missing GABA neurons (Fig. 3E-F). This tissue-specific effect demonstrates that the UNC-83a N-terminal domain specifically disrupts kinesin-1-mediated nuclear migration events, confirming its selective inhibitory function *in* vivo.

**Figure 3:**
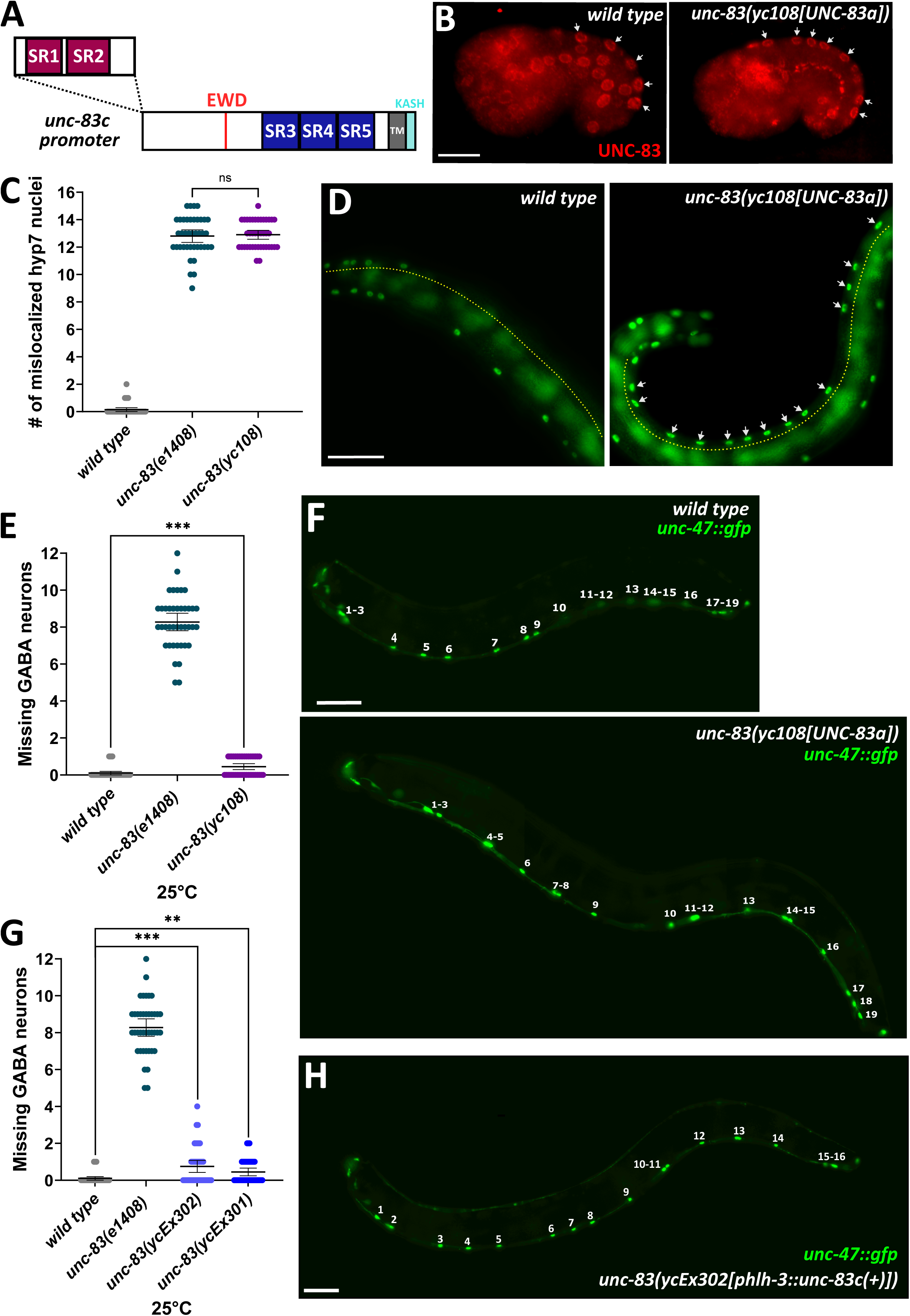
Ectopic expression of *unc-83a* and *unc-83c* disrupts kinesin-1-dependent hyp7 nuclear migration and dynein-mediated P-cell nuclear migration, respectively. A) Schematic of the *unc-83(yc108[UNC-83a])* mutant, expressing the UNC-83a isoform under control of the *unc-83c* promoter. B) UNC-83 (shown in red) in *unc-83(yc108[UNC-83a])* is localized to the nuclear envelope in a comma-stage embryo. Dorsal is up and anterior is left. Arrows point to the hyp7 nuclei. C) Quantification of hyp7 nuclear migration defects in wild type, *unc-83(yc108[UNC-83a])*, and *unc-83(e1408)* null animals via counting the number of mislocalized (at the dorsal cord) nuclei. Each point represents the total number of abnormally located hyp7 nuclei per animal; n = 40 for each strain. D) Lateral views of *wild type* and *unc-83(yc108[UNC-83a])* L4 larva expressing hypodermal nuclear GFP. Dashed yellow lines mark the dorsal cord of the animal. Mislocalized nuclei (arrows) indicate migration failure. E) Quantification of dynein-mediated larval P-cell nuclear migration defects in *wild type*, mutant *unc-83(yc108[UNC-83a])*, and *unc-83(e1408)* null animals via counting number of missing GABA neurons at 25°C. Each point represents the number of missing GABA neurons per animal; n = 40 for each strain. F) Lateral views of *wild type* and *unc-83(yc108[UNC-83a])* L4 larva expressing a GABA neuron nuclear marker (*unc-47::gfp*). Numbers indicate a total of 19 GABA neurons. G) Quantification of larval P-cell nuclear migration defects in wild type, *unc-83(e1408)* null, and *unc-83c* overexpression animals (*unc-83(ycEx301[phlh3::unc-83c(+)])* and *unc-83(ycEx302[phlh3::unc-83c(+)])*) via counting number of missing GABA neurons at 25°C. Each point represents the number of missing GABA neurons per animal; n = 40 for each strain. H) Lateral view of *unc-83(ycEx302)* L4 larvae expressing *unc-47::gfp*. Numbers indicate a total of 16 GABA neurons in larva. For all plots: Means with 95% CI are shown in error bars. Unpaired student t-tests were performed on the indicated comparisons; ns means not significant, p>0.05; * p<0.05; **** p<0.0001. For images: Scale bars = 10 μm for B), and 42.2 μm for D), F) and H).

### Ectopic expression of *unc-83c* disrupts dynein-mediated P-cell nuclear migration

Our findings suggested a model where UNC-83 isoforms establish motor protein bias through reciprocal inhibition. In this model, UNC-83a inhibits kinesin-1 while UNC-83c would be predicted to inhibit dynein. To test this prediction, we examined whether elevated UNC-83c levels in larval P cells could disrupt dynein-dependent P-cell nuclear migration. We generated transgenic animals ectopically overexpressing the short *unc-83c* isoform under control of the P-cell specific *hlh-3* promoter^63^. Two independent extrachromosomal arrays caused mild but significant P-cell nuclear migration defects with 0.45 + 0.20 and 0.75 + 0.33 missing GABA neurons compared to 0.1 ± 0.09 missing in wild-type animals (Fig. 3G, H). These mild defects support our model where ectopic UNC-83c spuriously activates kinesin-1 in P-cells, occasionally disrupting the normally dynein-dependent nuclear migration process.

### Spectrin-like repeats in the N-terminal domain of UNC-83a are necessary for dynein-dependent nuclear migration

We next sought to investigate the regulatory mechanism behind dynein-dependent P-cell nuclear migration. Previous genetic analysis revealed that mutations eliminating *unc*-*83a/b* while preserving *unc-83c* cause temperature-sensitive defects in P-cell nuclear migration^52^. To determine the critical regions within the UNC-83a/b N-terminus, we engineered a series of in-frame deletions targeting the predicted spectrin-like repeats (Fig. 4A). We confirmed that the deletion variants localized normally to the nuclear envelope by immunofluorescence (Fig. 4B). Quantitative analysis of P-cell nuclear migration revealed distinct requirements for different parts of the N-terminal domain. Disruption of both predicted spectrin-like repeats in *unc-83a(*Δ*58-233)* caused severe P-cell nuclear migration defects at 25°C (an average of 7.70 ± 0.75 missing GABA neurons), comparable to the *unc-83* null phenotype (Fig. 4C-D). Two smaller deletions in the spectrin-like repeat region, *unc-83a(*Δ*58-166)* and *unc-83a(*Δ*167-233)*, also led to null-like defects (Fig 4C). All three mutations exhibited temperature sensitivity, with defects progressively worsening from 15°C to 20°C and then to 25°C (Fig. 2S), matching the same temperature-sensitive phenotype of *unc-83* null animals. In contrast, a deletion just downstream of the predicted spectrin-like repeats, *unc-83a(*Δ*234-305)*, caused only minor P-cell nuclear migration defects (1.25 ± 0.45 missing GABA neurons) (Fig. 4C-D).

**Figure 4:**
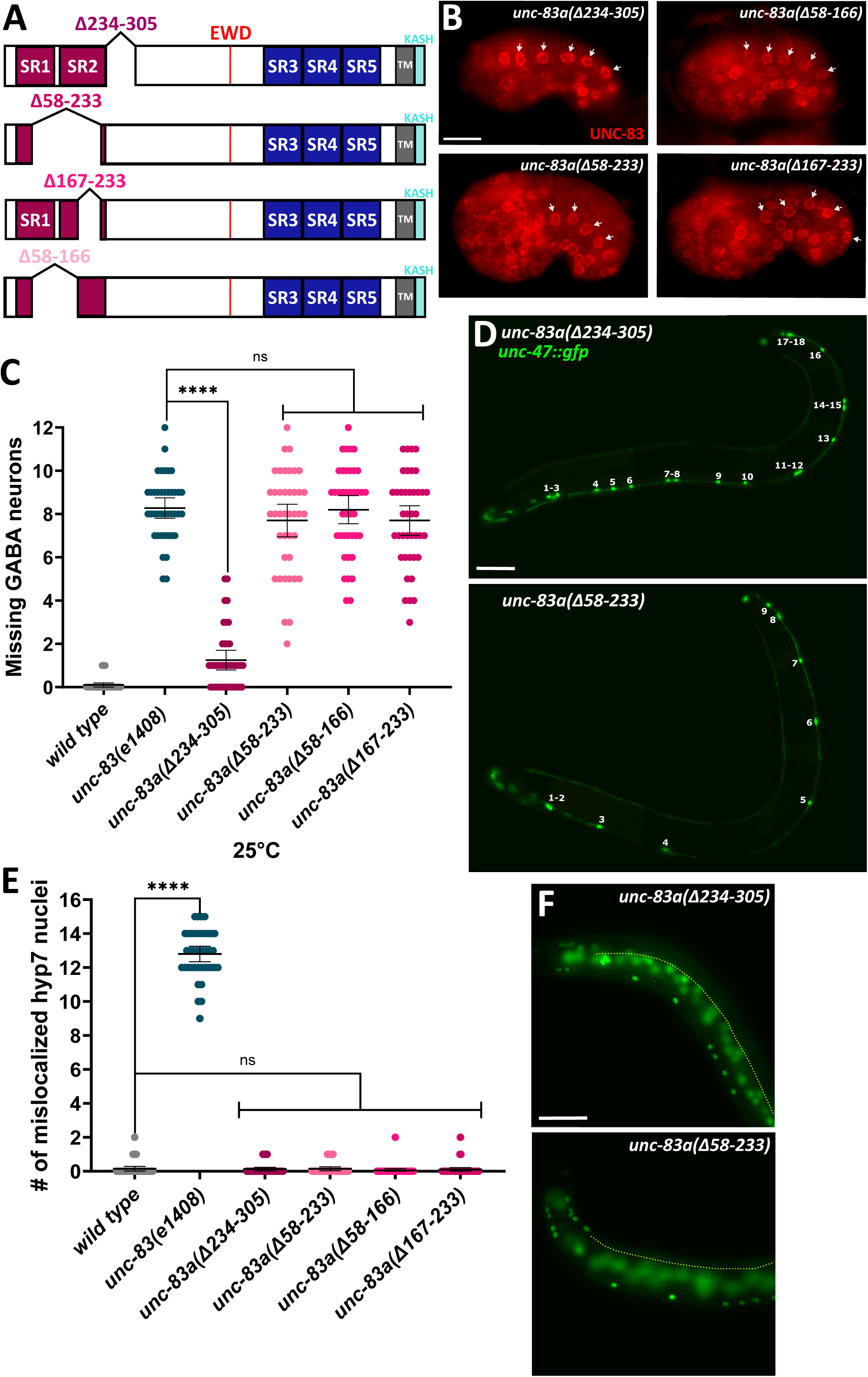
Spectrin-like repeats in the N-terminal domain of UNC-83a are necessary for dynein-dependent nuclear migration. A) Schematics of the *unc-83* N-terminal domain deletion mutants. B) Immunofluorescence staining shows that the deletion mutants *unc-83a(*Δ*234-305), unc*-*83a(*Δ*58-166)*, *unc-83a(*Δ*58-233)*, and *unc-83a(*Δ*167-233)* have a normal UNC-83 (red) localization pattern at the nuclear envelope of hyp7 precursor cell nuclei in comma-stage embryos. White arrows indicate nuclei with UNC-83 localization. Dorsal is up and anterior is left. C) Quantification of larval P-cell nuclear migration defects in the indicated strains at 25°C (See Figure S2 for the P-cell nuclear migration defects of the indicated strains at 20°C and 15°C.). Each point represents the number of missing GABA neurons per animal; n = 40 for each strain. D) Lateral views of *unc-83a(*Δ*234-305)* and *unc-83a(*Δ*58-233)* L4 larvae expressing *unc-47::gfp.* Numbers indicate total GABA neurons. E) Quantification of hyp7 nuclear migration defects in the indicated strains. Each point represents the total number of abnormally located hyp7 nuclei per animal; n = 40 for each strain. F) Lateral views of *unc-83a(*Δ*234-305)* and *unc-83a(*Δ*58-233)* L4 larvae expressing hypodermal nuclear GFP. Dashed yellow lines mark the dorsal cord of the animals. Arrows indicate mislocalized hypodermal nuclei, representing failed nuclear migrations. For all plots: Means with 95% CI are shown in error bars. Unpaired student t-tests were performed on the indicated comparisons; ns means not significant, p>0.05; * p<0.05; **** p<0.0001. For images: Scale bars = 10 μm for B) and 42.2 μm for D) and F).

None of these N-terminal deletions disrupted the role of *unc-83c* in embryonic hyp7 nuclear migrations. None had a significant accumulation of hyp7 nuclei in the dorsal cord (Fig. 4E-F). Together, these results demonstrate that the spectrin-like repeats in the UNC-83a/b N-terminal domain are required for dynein-dependent P-cell nuclear migration, but dispensable for kinesin-1-dependent hyp7 nuclear migration.

### A conserved enhancer controls cell-type specific expression of *unc-83c* in embryonic hypodermal precursors

The *unc-83* gene is hypothesized to have developmentally regulated expression of different isoforms, with *unc-83a/b* thought to be expressed in larval P cells and *unc-83c* in embryonic hyp7 precursors. We identified potential enhancer elements within the unusually long introns just upstream of the first *unc-83c* exon. Comparative genomic analysis across four *Caenorhabditis* species (*C. elegans*, *C. brenneri*, *C. remanei* and *C. briggsae*) identified a highly conserved 870bp sequence (Fig. 3S; Fig. 5A).

**Figure 5:**
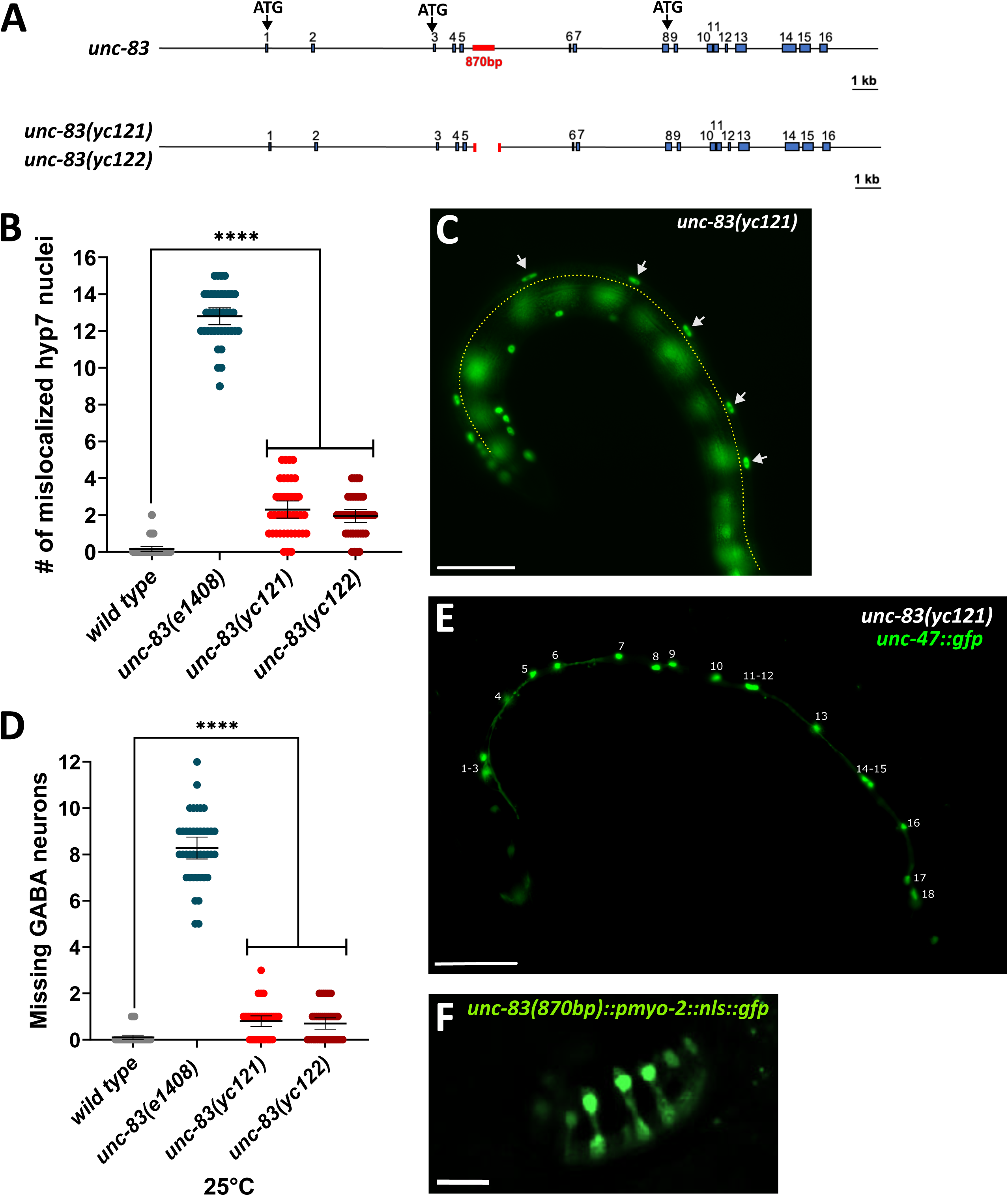
A conserved enhancer controls cell-type specific expression of *unc-83c* in embryonic hypodermal precursors. A) Schematics of the *unc-83* locus in *wild type,* and enhancer deletion mutants (*unc-83(yc121)* and *unc-83(yc122)*). The 870bp enhancer region is highlighted in red. Exons are shown as blue boxes. The start codon for *unc-83a* in exon 1, *unc-83b* in exon 3, and *unc-83c* at the end of exon 8 are indicated. The sizes of introns and exons are according to the scale as indicated (scale bar=1 kb). (See Figure S3 for the sequence alignment). B) Quantification of hypodermal hyp7 nuclear migration defects in the indicated strains. Each point represents the total number of abnormally located hyp7 nuclei per animal; n = 40 for each strain. C) Lateral view of *unc-83(yc121)* L4 larva expressing hypodermal nuclear GFP. Dashed yellow line mark the dorsal of the animal. Arrows indicate mislocalized hypodermal nuclei, representing failed nuclear migrations. D) Quantification of larval P-cell nuclear migration defects in the indicated strains at 25°C. Each point represents the number of missing GABA neurons per animal; n = 40 for each strain. E) Lateral view of *unc-83(yc121*) L4 larva expressing *unc-47::gfp*. Numbers indicate a total number of 18 GABA neurons. F-G) *C. elegans* pre-comma stage embryos expressing *unc-83* 870bp enhancer region driving *nls::gfp*. F) Expression of the 870 bp enhancer region (green) in pre-comma stage embryo, showing hyp7 precursor-specific expression. For all plots: Means with 95% CI are shown in error bars. Unpaired student t-tests were performed on the indicated comparisons; ns means not significant, p>0.05; * p<0.05; **** p<0.0001. For images: Scale bars = 42.2 μm for C) and E) and 10 μm for F) and G).

To test the necessity of this predicted enhancer, we generated two independent alleles, *unc-83(yc121)* and *unc-83(yc122)*, where we deleted 750bp of this region using CRISPR/Cas9 gene editing (Fig. 5A). Quantitative analysis of embryonic hyp7 and larval P-cell nuclear migrations revealed identical phenotypes in *unc-83(yc121)* and *unc-83(yc122)* strains. An average of about two hyp7 nuclei mislocalized in the dorsal cord (Fig. 5B-C), significantly higher than in wild type, indicating the enhancer’s necessity for hyp7 nuclear migration. The enhancer deletion also caused subtle but significant defects in P-cell nuclear migration (Fig. 5D-E).

To test the enhancer’s sufficiency to drive the expression of *unc-83c* in embryonic hyp7 precursors, we generated a transgene by fusing the 870bp region to a minimal promoter driving the expression of GFP fused to the SV40 nuclear localization sequence (*nls::gfp*). This construct specifically drove expression in pre-comma stage embryonic hyp7 precursors (Fig. 5F). These results demonstrate that this conserved non-coding 870bp element functions as a cell-type specific enhancer driving *unc-83c* expression in embryonic hyp7 precursors. This is consistent with the earliest detected expression pattern of UNC-83 protein^52^.

### The UNC-83a N-terminal domain inhibits kinesin-1 activity *in vitro*

Building on our genetic evidence that the UNC-83a-specific N-terminal domain inhibits kinesin-1 while promoting dynein activity during P-cell nuclear migration, we investigated its direct effects on kinesin-1 activity using biochemical approaches. We expressed and purified the N-terminal domain of UNC-83a (UNC-83a^N-term^) from *E. coli* (Fig. 6A-B). Additionally, we purified recombinant *C. elegans* kinesin-1 heterotetramers from Sf9 insect cells co-expressing full-length KHC (UNC-116) and KLC (KLC-2) (Fig. 6A-B). As mammalian kinesin-1 heterotetramers require cargo adaptors for activation from their autoinhibited state^30,64^, we first analyzed *C. elegans* kinesin-1 recruitment to microtubules in the presence of a non-hydrolysable nucleotide analog, AMP-PNP, which locks the motor domains in a high-affinity state on microtubules^64^ (Fig. 6C-D). Kinesin-1 alone bound to microtubules as expected^64^, whereas neither UNC-83a nor UNC-83c showed any observable microtubule binding in the absence of kinesin (Fig. 6C). Addition of excess UNC-83a or UNC-83c significantly enhanced kinesin-1 binding to microtubules, with UNC-83c demonstrating greater efficiency (Fig. 6C-D). In contrast, UNC-83a^N-term^ failed to increase kinesin-1 recruitment to microtubules above baseline levels (Fig. 6C-D).

**Figure 6:**
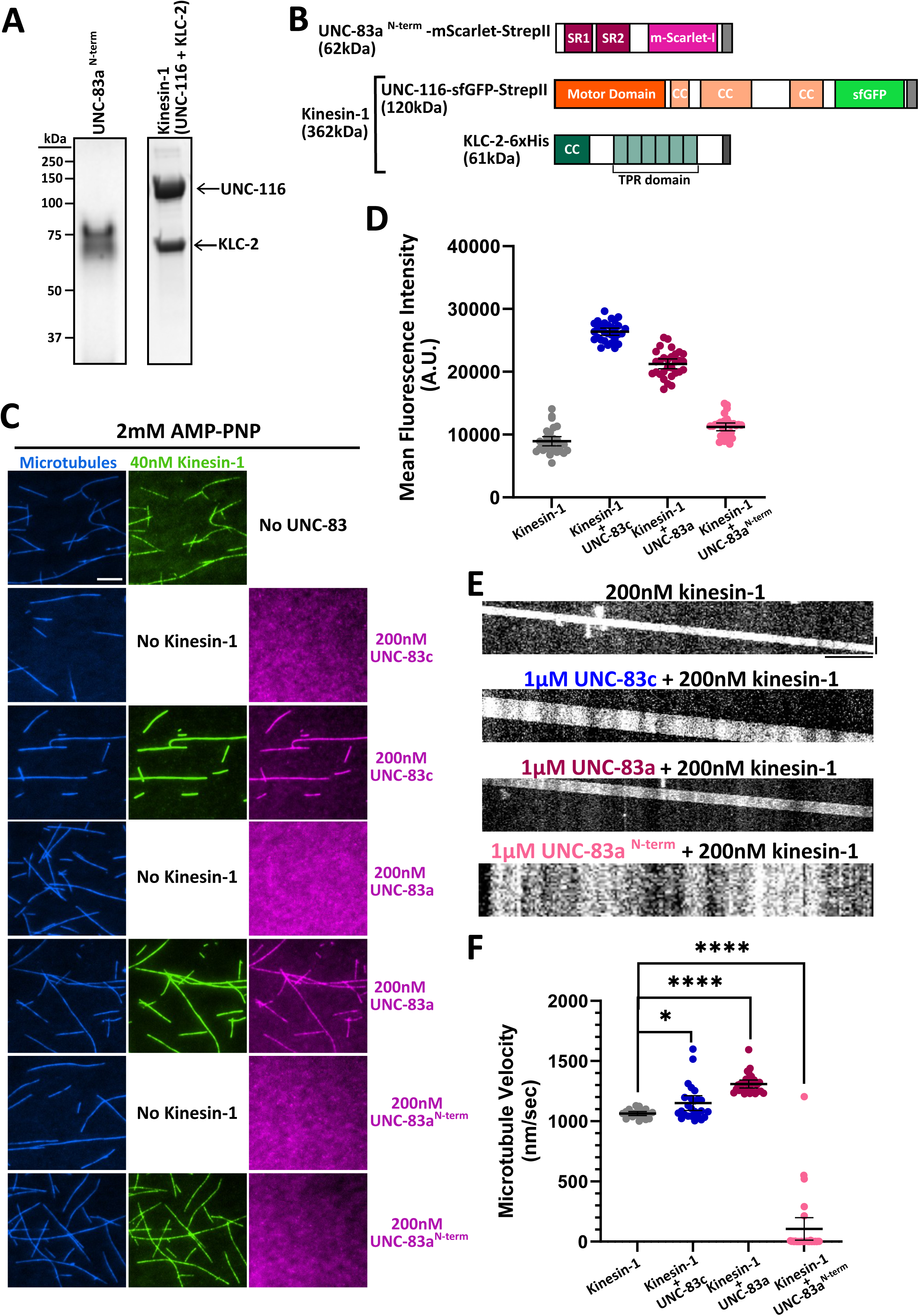
The UNC-83a N-terminal domain inhibits kinesin-1 activity *in vitro*. A) Coomassie-stained gels of purified UNC-83a^N-term^ and UNC-116-KLC-2 (kinesin-1) recombinant proteins. Arrows indicate the bands for the different proteins. B) Schematics of the indicated recombinant protein constructs. The coiled-coil domains of UNC-116 and KLC-2 are indicated as ‘CC’. UNC-116 is co-expressed with KLC-2. The co-expression leads to the formation of kinesin-1 heterotetramer that is expected to be 362 kDa. C) Representative TIRF images showing the recruitment of kinesin-1 (green) in the rigor state (with 2 mM AMP-PNP) onto microtubules (blue) and the effects of the indicated UNC-83 constructs (magenta). Scale bar = 5 µm. D) Quantification of fluorescence intensity the indicated constructs, n = 30 microtubules for each condition. E) Representative kymographs of microtubule gliding with kinesin-1 alone (See Movie S1), kinesin-1 + UNC-83c (See Movie S2), kinesin-1 + UNC-83a (See Movie S3), and kinesin-1 + UNC-83a^N-term^ (See Movie S4). Scale bars: 20 s (vertical) and 5 µm (horizontal). F) Quantification of microtubule gliding velocities for the indicated conditions, n = 25 microtubules for each condition. The average velocities are: 1,064 ± 35 nm/s (kinesin-1), 1,150 ± 149 nm/s (kinesin-1 + UNC-83c), 1,309 ± 82 nm/s (kinesin-1 + UNC-83a), 187 ± 533 nm/s (kinesin-1 + UNC-83a^N-term^).

We next assessed the impact of UNC-83a, UNC-83c and UNC-83a^N-term^, on kinesin-1 motor function using microtubule gliding assays (Fig. 6E-F, Movie S1-4). Both UNC-83a and UNC-83c enhanced microtubule gliding velocity when added in excess to kinesin-1 (Movie S2-3). Strikingly, UNC-83a^N-term^ significantly reduced the average gliding velocity compared to kinesin-1 alone (Figs. 6E-F and Movies S1 and S4), providing direct evidence that the N-terminal domain of UNC-83a functions as a kinesin-1 inhibitor. Importantly, we saw no evidence of UNC-83a or UNC-83a^N-term^ binding directly to microtubules (Fig. 6C), indicating the blockage of kinesin-1 motility is a direct effect on kinesin and not a secondary effect of microtubule-binding.

### The UNC-83a N-terminal domain directly binds kinesin heavy chain

Having established that UNC-83a inhibits kinesin-1 activity both *in vivo* and *in vitro*, we investigated the molecular mechanism underlying this inhibition. We hypothesized that the UNC-83a^N-term^ domain inhibits kinesin-1 by directly binding the KHC UNC-116. To test this hypothesis, we purified full-length UNC-116 from Sf9 cells (Fig. 7A), confirming that the purified protein existed predominantly (82%) in its expected dimeric state^28^ (Fig. 7B). We then performed binding studies by separately incubating UNC-83a^N-term^, UNC-83a, and UNC-83c with UNC-116 (Fig. 7D’-F’). Both UNC-83a^N-term^ and UNC-83a formed stable complexes with UNC-116 (one monomer of UNC-83a or UNC-83a^N-term^ binding one dimer of UNC-116) as detected by mass photometry (Fig. 7D-D’, E-E’). In contrast, UNC-83c failed to interact with UNC-116 (Fig. 7F-F’). We then performed binding studies by separately incubating UNC-83a and UNC-83c with kinesin-1 tetramer (UNC-116 + KLC-2) (Fig. 7C). We observed one UNC-83a monomer forming a stable complex with one kinesin-1 tetramer while UNC-83c did not (Fig. 7D’’, F’’). These findings demonstrate that the UNC-83a-specific N-terminal domain mediates direct binding to UNC-116, providing a molecular mechanism for the observed kinesin-1 inhibition both *in vivo* (Fig. 3A-D) and *in vitro* (Fig. 6E-F).

**Figure 7:**
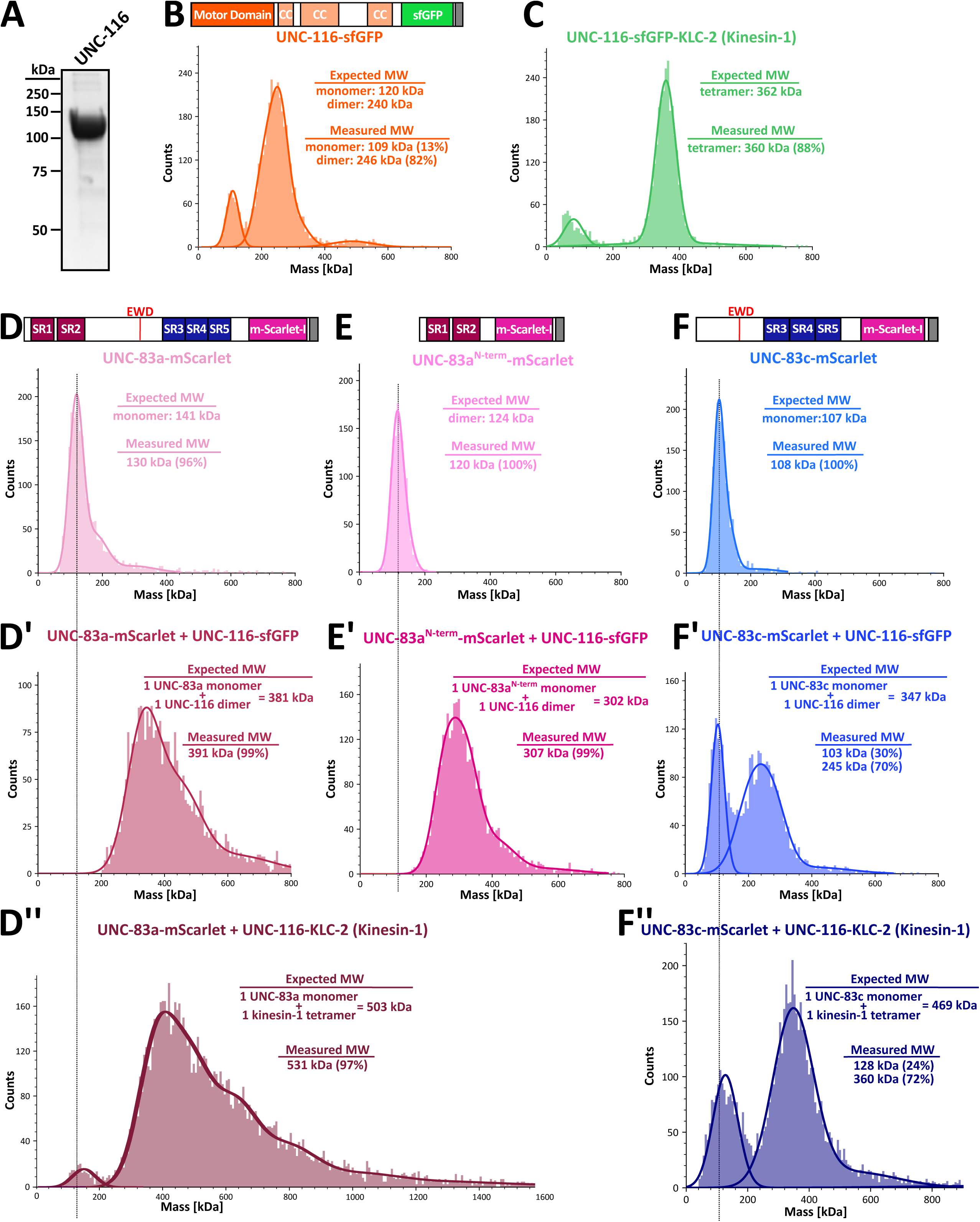
The UNC-83a N-terminal domain directly binds kinesin heavy chain. A) A Coomassie-stained gel showing purified UNC-116-sfGFP. B) A schematic of the UNC-116-sfGFP recombinant protein and a Gaussian fit of its measured molecular weight in solution by mass photometry (30 nM; monomer = 109 ± 19.5 kDa, 13% peak; dimer = 246 ± 44 kDa; 82% peak). Coiled-coil domains indicated as ‘CC’. C) A Gaussian fit of UNC-116-KLC-2 (kinesin-1) recombinant protein’s measured molecular weight in solution by mass photometry (30 nM; tetramer = 360 ± 69 kDa, 88% peak). D-D’’) A schematic of the UNC-83a-mScarlet recombinant protein and a Gaussian fit of its measured molecular weight in solution alone (30 nM; monomer = 130 ± 62 kDa; 96% peak) (D), when preincubated with UNC-116 (30 nM; one UNC-83a monomer + one UNC-116 dimer = 391 ± 153 kDa, 99% peak) (D’), and when preincubated with UNC-116-KLC-2 (kinesin-1) (30 nM; one UNC-83a monomer + one kinesin-1 tetramer = 531 ± 272 kDa, 97% peak) (D’’). E-E’) A schematic of the UNC-83a^N-term^-mScarlet recombinant protein and a Gaussian fit of its measured molecular weight in solution alone (30 nM; dimer = 120 ± 23 kDa, 100% peak) (E) or when preincubated with UNC-116 (30 nM; one UNC-83a^N-term^ monomer + one UNC-116 dimer = 307 ± 89 kDa, 99% peak) (E’). F-F’’) A schematic of the UNC-83c-mScarlet recombinant protein and a Gaussian fit of its measured molecular weight in solution alone (30 nM; monomer = 108 ± 41 kDa, 100% peak) (F), when preincubated with UNC-116 (30 nM; 103 ± 21 kDa, 30%; 245 ± 82 kDa, 70% peak) (F’), and when preincubated with UNC-116-KLC-2 (kinesin-1) (30 nM; 128 ± 41 kDa, 24% peak; 360 ± 110 kDa, 72% peak) (F’’).

## DISCUSSION

Spatial and temporal control of nuclear positioning is fundamental for development, yet the mechanisms regulating bidirectional nuclear movement have remained elusive. Our work reveals a novel paradigm where expression of alternative isoforms of a cargo adaptor creates an intrinsic molecular switch for controlling nuclear migration directionality. Previous studies identified UNC-83 as a nuclear-specific cargo adaptor that recruits both kinesin-1 and dynein to mediate bidirectional movement of nuclei^15,55,56^. Genetic analyses suggested a model where the short UNC-83c isoform is sufficient for kinesin-driven nuclear migration in the embryo and the long UNC-83a/b isoforms are necessary for dynein-mediated nuclear migration in larva^52^. Here we demonstrate that distinct UNC-83 isoforms differentially regulate motor activity to achieve precise developmental control of nuclear positioning.

Our discovery of an inhibitory N-terminal-specific domain in the long UNC-83a/b isoforms represents a previously unrecognized mechanism for motor regulation. The genetic and biochemical analyses reported here reveal this domain’s dual regulatory function: direct binding and inhibition of the kinesin heavy chain UNC-116, combined with reduced affinity for the kinesin light chain KLC-2. This regulation allows UNC-83a/b to suppress kinesin-1 activity while permitting dynein-directed movement. In contrast, UNC-83c’s high-affinity interaction with KLC-2 through its EWD motif^13,36^ promotes kinesin-1 activation, enabling plus-end-directed nuclear movement in embryonic hyp7 precursors. This isoform-specific regulation solves a challenge in developmental biology: how to achieve opposing directional movements of the same cargo using the same molecular machinery.

The involvement of UNC-83 with multiple dynein-regulatory proteins, including the homolog of long coiled-coil dynein activating adaptor Bicaudal D, BICD-1^56^, suggests integration into a larger regulatory network. This organization parallels other bidirectional transport events^65,66^. For example, TRAK1/2 mediates mitochondrial trafficking through the outer mitochondrial membrane receptor Miro and metaxins^67,68,69^, and HOOK3 regulates endosomal trafficking with FTS and FHIP1B^70,71,72,73^. Nuclei, which are larger and stiffer than mitochondria and endosomal vesicles^74^, present unique challenges for intracellular transport. Our findings and the conservation of LINC complex-mediated nuclear positioning^17,22,54^ suggest that evolution has developed specialized mechanisms to address the challenges of moving nuclei. Mammalian KASH proteins Nesprin-2, Nesprin-4, and KASH5 each interact with microtubule motors through mechanisms similar to UNC-83^13,14,60,41,42,64,75^.

The identification of spectrin-like repeats in UNC-83 with AlphaFold predictions reveals structural similarities to *Drosophila* Klarsicht^76^ and mammalian nesprins^13,14,60,41,42,64,75^. This structural conservation, combined with common motor-binding mechanisms, suggests that motor regulation at the nuclear envelope through alternatively spliced inhibitory domains may be more widespread than previously appreciated. The modular nature of KASH protein interactions with the cytoskeleton appears to be an evolutionarily conserved feature, potentially allowing rapid adaptation to diverse cellular contexts.

Our data support a selective activation model for bi-directional transport, rather than selective recruitment or tug-of-war models^66^. The selective activation model could explain the steady, but slow nature of nuclear movements^77^. Moreover, cargo stiffness has been speculated to alter motor engagement^78,79^. Nuclei are the largest and one of the stiffer intracellular cargos and move significantly slower than small cargos. Hyp7 and P-cell nuclei migrate at a speed of ∼1µm per minute^49,56^. The discovery that expression of alternative isoforms can generate cargo adaptors with opposing regulatory functions provides a new framework for understanding how cells achieve precise spatiotemporal control of organelle positioning. This mechanism may be particularly crucial for nuclei, where their unique physical properties^74^ require additional regulation.

These findings open several exciting directions for future investigation. Does Klarsicht employ similar inhibitory mechanisms during *Drosophila* development? Have mammalian systems evolved analogous regulatory strategies through their expanded repertoire of KASH proteins? Understanding how broadly this regulatory paradigm extends could reveal fundamental principles of developmental organelle positioning. Moreover, our work suggests that expression of alternative isoforms of cargo adaptors may represent a general mechanism for generating precise spatiotemporal control of intracellular transport during development.

## RESOURCE AVAILABILITY

### Lead contact

Further information and requests for resources and reagents should be directed to and will be fulfilled by the lead contact, Daniel A. Starr

### Materials availability

All unique reagents generated in this study are available from the lead contact without restriction.

### Data and code availability

Data reported in this paper will be shared by the lead contact upon request. This paper does not report original code. Any additional information required to reanalyze the data reported in this paper is available from the lead contact upon request.

## ACKNOWLEDGEMENTS

We thank Alyssa Paparella and Maryam Alani (UC Davis) for their technical assistance in *C. elegans* genetics. We thank Gino Cortopassi and Alexey Tomilov (UC Davis) for assistance with the Octet RED 385 instrument. We thank Regina Bohn, Eve Ladwig, and Esther Oh (UC Davis) for their early contributions to the *unc-83* enhancer work. We thank WormBase for its valuable resources. This research was supported by the National Institutes of Health and the NIGMS through grants 1R35GM124889 to RJM and R35GM134859 to DAS.

## AUTHOR CONTRIBUTIONS

Conceptualization: SG, GWGL, RJM, DAS. Data curation: SG, NS, DE, EFG, KC. Formal analysis: SG, NS, DE, EFG, KC. Investigation: SG, NS, DE, EFG, KC. Methodology: all authors. Project Administration: SG, DAS. Resources: all authors. Validation: all authors. Visualization: SG, GWGL, RJM, DAS. Writing-original draft: SG, DAS. Writing-review and editing: SG, SN, GWGL, RJM, DAS.

## EXPERIMENTAL MODEL AND SUBJECT DETAILS

### *C. elegans* maintenance and genetics

*C. elegans* strains were maintained at room temperature (approximately 22°C), unless otherwise indicated, on nematode growth medium plates seeded with OP50 *E. coli* ^80^. Some strains were obtained from the Caenorhabditis Genetics Center, funded by the National Institutes of Health Office of Research Infrastructure Programs (P40 OD010440). All the strains used in this study are listed on the Key Resources Table.

### Protein expression systems

*Spodoptera frugiperda* Sf9 cells were cultured in suspension using Sf-900II serum-free medium (SFM) (ThermoFisher Scientific) supplemented with 10 µg/mL gentamycin. Cultures were maintained at 27°C with constant rotation at 150 rpm. The culture density was maintained between 1.0×10^6^ and 8.0×10^6^ cells/mL during routine passaging. Cells were not authenticated. The sex of the cells is unknown.

BL21-CodonPlus (DE3)-RIPL *E. coli* (Agilent) transformed with expression plasmids were cultured in LB medium at 37°C with shaking at 200 rpm.

## METHOD DETAILS

### Protein structure prediction

Structural models of UNC-83a and the UNC-83c/KLC-2 complex were generated using AlphaFold3^59^ with default parameters, without template structures. Five models were generated for each protein or complex. The models with the highest confidence scores (for the UNC-83c/KLC-2 complex: pTM = 0.44 and ipTM = 0.39) were used for figure preparation. For visualization, some residues within the low-confidence unstructured region between the spectrin-like repeats #3 and #4 in UNC-83a were straightened by altering the torsion angles^81^ (φ and ψ to approximately –135° and 135°, respectively) with ChimeraX^82^.

### Plasmid construction and molecular cloning

#### Expression constructs

The glutathione-S-transferase (GST)-KLC-2 construct was generated as previously described^55^. A cDNA construct encoding for UNC-83a^52^ was used to generate the UNC-83a-mScarlet-StrepII, UNC-83c-mScarlet-StrepII, and UNC-83a^N-term^-mScarlet-StrepII constructs. Using Gibson Assembly, UNC-83a, UNC-83c and UNC-83a^N-term^ coding regions were cloned into pET21a-Novagen bacterial expression vector (Millipore Sigma) with a C-terminal mScarlet-StrepII tag. For insect cell expression, UNC-116-sfGFP-StrepII and KLC-2-6×His-UNC-116-sfGFP-StrepII constructs were cloned into pACEBac1 vector (Geneva Biotech).

#### Promoter and enhancer constructs

The pSL536 plasmid was made by cutting the 3.4 kb fragment of *hlh-3* promoter from pRD15^63^ (using HindIII and SmaI restriction enzyme sites) and ligating this fragment 442 bp upstream of the *unc-83c* start codon into AgeI/BamHI/Klenow treated pDS22^52^. For pSL439, the 870bp intronic fragment from the *unc-83* region between exons 5 and 6 was cloned using StuI and XbaI, and inserted into pPD97.82 (Addgene plasmid #1512, a gift from the Fire Lab). The *odr-1::rfp* plasmid was a gift from Noelle L’Etoile (University of California, San Francisco)^83^.

#### Sequence verification

All constructs were verified by whole plasmid sequencing performed by Plasmidsaurus using Oxford Nanopore Technology with custom analysis and annotation. A complete list of plasmids used in this study is provided in the Key Resources Table.

### Protein expression and purification

Bacterial expression and preparation of GST-KLC-2, UNC-83a-mScarlet-StrepII, UNC-83c-mScarlet-StrepII, and UNC-83a^N-term^-mScarlet-StrepII were performed as described below. BL21-CodonPlus (DE3)-RIPL (Agilent) *E. coli* were transformed with each plasmid and grown at 37°C in LB medium with 100 µg/mL carbenicillin until optical density at 600 nm (OD_600_) reached 0.8. The cultures were then cooled down to room temperature and protein expression was induced by adding 0.2 mM isopropyl-b-D-thiogalactoside (IPTG) and incubating overnight at 18°C. Bacterial pellets were stored at –80°C. For recombinant protein purification, frozen pellets were thawed on ice and resuspended in 50 mL lysis buffer (PB: 50 mM Tris-HCl pH 8.0, 150 mM KCH3COO, 2 mM MgSO4, 1 mM EGTA, and 5% glycerol) freshly supplemented with 1 mM dithiothreitol (DTT), 0.2 mM phenylmethyl-sulfonyl fluoride (PMSF), 0.1 mM ATP, NucA nuclease, and protease inhibitor mix (Promega). Bacteria were lysed using an Emulsiflex C3 high-pressure homogenizer (Avestin). For UNC-83a-mScarlet-StrepII, UNC-83c-mScarlet-StrepII, and UNC-83a^N-term^-mScarlet-StrepII purifications, a subsequent chemical lysis was performed with 0.15% Triton X-100 for 5 minutes on ice. Lysed cells were centrifuged at 23,000xg for 20 minutes at 4°C. For affinity purification, the clarified lysates were incubated with resin as follows. UNC-83a-mScarlet-StrepII, UNC-83c-mScarlet-StrepII, and UNC-83a^N-term^-mScarlet-StrepII lysates were incubated with Streptactin XT resin (IBA Lifesciences) for 1 hour at 4°C and washed with PB buffer. Proteins were eluted with biotin elution buffer (100mM biotin in PB buffer freshly supplemented with 1 mM DTT, 0.1 mM PMSF, 0.1 mM ATP, NucA nuclease, 0.013% Triton X-100 at pH 8.0). For GST-KLC-2, Glutathione Sepharose 4B resin (Cytiva) was added to the clarified lysate and incubated on a rocker for 5 hours at 4°C. The beads were then washed with 1x PBS (Phosphate-buffered saline) and protein was eluted with glutathione elution buffer (50 mM Tris-HCl pH8.0, 10 mM L-Glutathione reduced; (Sigma-Aldrich).

For the UNC-116-sfGFP-StrepII and KLC-2-6xHis-UNC-116-sfGFP-StrepII constructs cloned into pACEBac1 vector, DH10MultiBac (Geneva Biotech) *E. coli* were transformed to generate bacmids. To prepare baculovirus, 1.0×10^6^ Sf9 cells were transferred to each well of a tissue-culture treated 6-well plate. Once the cells were adhered to the bottom of the dish, about 3 µg bacmid were transfected using 6 µL of Cellfectin II reagent (ThermoFisher Scientific). 5-7 days after initial transfection, the culture media were collected and spun at 3,000xg for 3 minutes to obtain the viral supernatant (P1). Next, 50 mL of Sf9 cells (2.0 x10^6^ cells/mL) was infected with 30 µL of P1 and cultured for 5-7 days to obtain P2 viral supernatant. For protein expression, 400 mL of Sf9 cells (2.0×10^6^ cells/mL) were infected with 4 mL of P2 virus and cultured for 65 hours at 27°C while shaking. Cells were harvested by spinning down at 1,992xg for 20 minutes at 4°C. The pellets were stored at –80°C.

UNC-83a^N-term^-mScarlet-StrepII, UNC-116-sfGFP-StrepII and KLC-2-6xHis-UNC-116-sfGFP-StrepII were further purified via size exclusion chromatography using Superose6 10/300 GL column (Cytiva) equilibrated in buffer GF150 (25 mM HEPES pH 7.2, 150 mM KCl, 2 mM MgCl_2_). GST-KLC-2, UNC-83a-mScarlet-StrepII and UNC-83c-mScarlet-StrepII were purified via size exclusion chromatography using Superdex 200 10/300 GL column (Cytiva) equilibrated in buffer GF150. Fractions containing the proteins were combined and concentrated on Amicon spin filters with a 50 kDa or 100 kDa cutoff. Concentrated proteins were supplemented with 10% glycerol, flash frozen in LiN2 and stored at C-80°C.

### Single-molecule mass photometry

Single-molecule mass photometry was performed using a OneMP instrument (Refeyn) as previously described^60, 84^. Briefly, #1.5 24 x 50 mm Deckgläser glass coverslips (ThermoFisher Scientific) were cleaned by 1 hour sonication in isopropanol and subsequently stored in a fresh aliquot of isopropanol. Before assembling the imaging chambers, the coverslips were rinsed with fresh isopropanol followed by Milli-Q water and dried with filtered air. Then, CultureWell silicone gaskets (Grace Bio-Labs) were placed onto the freshly cleaned coverslips providing 4-6 independent sample chambers. For all the cleaning steps, 99.9% (HPLC grade) isopropanol (Fisher Scientific) was used. Samples and standard proteins (Apoferritin, Bovine serum albumin (BSA), and Thyroglobulin; Sigma-Aldrich) were diluted to 10-100 nM in freshly 0.22 μm filtered SRP90 buffer (90 mM HEPES-KOH, 50 mM KCH3COO, 2 mM Mg(CH3COO)2, 1 mM EGTA, and 10% glycerol, pH = 7.6). The focal position was identified and maintained throughout the measurement using an autofocus system based on total internal reflection. Data acquisition was performed at 1 kHz for 60 seconds using AcquireMP (Refeyn), and data analysis was conducted using DiscoverMP (Refeyn).

### Biolayer interferometry binding assays

The real-time binding kinetics of UNC-83a-mScarlet-StrepII and UNC-83c-mScarlet-StrepII to GST-KLC-2 were measured using biolayer interferometry on a ForteBio Octet^®^-RED384 at the UC Davis Octet^®^ Real-Time Drug and Protein Binding Kinetics Unit. Binding assays were performed in binding buffer (20 mM Tris pH7.8, 150 mM NaCl, 0.1 mg/mL BSA, 0.05% Tween-20) (modified from Bhattacharya et al., 2014)^85^ using the Octet^®^ Anti-Glutathione-S-Transferase (GST) Biosensors (Sartorius AG). Operating temperature was maintained at 26°C with a 1,000 rpm rotary agitation throughout the experiment. Samples or buffer were transferred into 96-well plates (Greiner) at a volume of 250 μL per well. GST biosensor tips were rehydrated in binding buffer before loading the bait onto them. The bait, GST-KLC-2, was diluted to 1 µM and immobilized onto the biosensors for 10 minutes to attain 1.5 – 2.5 nm response. Binding association of UNC-83a and UNC-83c with biosensor tips was monitored for 20 minutes, and subsequent disassociation in buffer was monitored for 30 minutes. Both UNC-83a and UNC-83c were tested at concentrations of 1.0, 0.667, 0.444, 0.296, 0.197, 0.132, 0.088 μM. The average of the last 100 points (20 seconds) of the of the association phase response was used as an estimate for the equilibrium value of the binding reaction. The plots of concentration versus averaged plateau values were fitted using a quadratic binding equation plus a constant offset to find K_d_.

### *C. elegans* transgenic strains

To generate transgenic strains UD336 and UD338, which express nuclear GFP (*nls::*GFP) under a minimal promoter with the 870 bp predicted *unc-83* enhancer, N2 animals were injected with 5 ng/µL pSL439 and 100 ng/µL *odr-1::rfp* co-injection marker^83^. The integrated transgene *ycIs3[unc-83(870bp)::min_myo-2 promoter::nls::gfp; odr-1::rfp]* was generated with gamma irradiation from a Cs-137 source^86^. For strains UD973 and UD974, *oxIs12* (EG1285) animals were injected with 5 ng/µl of pSL536 (*phlh-3::unc-83c*), 93 ng/µl of pBlue-script SK, and 2 ng/µl of Pmyo-2::mRFP.

*unc-83a* deletion mutant alleles (*yc77,yc79, yc95* and *yc123*) and *unc-83* enhancer region deletion mutant alleles (*yc121* and *yc122*) were generated using the *dpy-10* co-CRISPR strategy^87, 88, 89^. The CRISPR injection mix was made as described previously^90^. As repair templates, 100-nucleotide-long single-strand DNA oligonucleotides (ssODN) were synthesized (Integrated DNA Technologies). Deletion mutations were screened by PCR amplification of the guide target region, and products showing a smaller band size were confirmed via Sanger sequencing. The crRNA and ssODN repair template sequences are listed in the Key Resources Table. *unc-83a^N-term^* overexpression insertion mutant allele (yc108) was generated by using *zen-4(+)* as a co-CRISPR marker^91^. A single-stranded repair template containing the insertion flanked by 50-nucleotide right and left homology arms, was synthesized (Genewiz). The CRISPR injection mix contained 0.2 μl *zen-4* crRNA (0.6 mM), 0.5 μl target gene crRNA (0.6 mM), 2.46 μl tracr (0.17 mM), 7.68 μl Cas9 (40 μM), 0.28 μl ssODN *zen-4(+)* repair template (500 ng/μl) and 2 μl ssDNA repair template (1µg/μl). The mixture was injected into the germline of temperature-sensitive *zen-4(cle10)* mutant young adults and screened as previously described^91^.

### Nuclear migration assays in *C. elegans*

To quantify larval P-cell nuclear migration, strains were crossed into *oxIs12[punc-47::gfp],* which labels P-cell derived GABA neurons^92^. The presence of 19 GABA neurons (12 derived from P cells) that are normally GFP-positive with cell bodies located in the ventral cord were scored for as previously described^62^. The assay was performed at 3 different temperatures (15°C, 20°C and 25°C) as *unc-83-*dependent P-cell nuclear migration is temperature-sensitive^52^. To quantify embryonic hyp7 nuclear migration, strains were crossed into *ycIs9 I,* a GFP-expressing nuclear marker specific to larval hypodermal nuclei^20^. Syncytial hyp7 nuclei were scored as abnormally located if they were in the dorsal cord as previously described ^62^.

### Immunofluorescence in *C. elegans*

Immunofluorescence staining with the anti-UNC-83 antibody (1209D7) was done as previously described^52^. The primary antibody 1209D7 was used undiluted. The secondary antibody, Alexa Fluor® 555 conjugated goat anti-mouse (H+L) Cross-Adsorbed IgG (ThermoFisher Scientific) was diluted 1:200 in phosphate-buffered saline containing 0.1% Triton X-100 (PBST). DNA was visualized by a 10-minute stain in 100 ng/ml of 4,6-diamidino-2-phenylindole (DAPI; Sigma) in PBST. Images were captured using a Leica DM6000 fluorescence microscope with a 63 Å∼ Plan Apo 1.40 NA objective, a Leica DC350 FX camera, and Leica LAS AF software.

### Microtubule preparation

Porcine brain tubulin was isolated using the high-molarity PIPES procedure and then labeled with biotin NHS ester or Dylight-405 NHS ester (ThermoFisher Scientific), as described previously (https://mitchison.hms.harvard.edu/files/mitchisonlab/files/labeling_tubulin_and_quantifying_lab eling_stoichiometry.pdf). Pig brains were obtained from a local abattoir and used within ∼4 hours. post-mortem. To polymerize microtubules, 50 mΜ of unlabeled tubulin, 10 μM of biotin-labeled tubulin and 3.5 μM of Dylight-405-labeled tubulin were incubated with 2 mM GTP for 20 minutes at 37°C. Polymerized microtubules were stabilized by the adding 20 μΜ taxol (ThermoFisher Scientific) and incubated an additional 20 minutes at room temperature. Microtubules were pelleted at 20,000 x g by centrifugation through a 150 μl of 25% sucrose cushion. The pellet was resuspended in 50 μl BRB80 (80 mM Pipes (pH 6.8), 1 mM MgCl2 and 1 mM EGTA) containing 10 μΜ taxol.

### TIRF assays

For TIRF assays, imaging chambers were assembled by combining an acid-washed glass coverslip prepared as previously described (http://labs.bio.unc.edu/Salmon/protocolscoverslip-preps.html), a precleaned slide, and double-sided sticky tape. Microtubules were diluted in BRB80 containing 10 μΜ taxol. SRP90 assay buffer (90 mM HEPES, 50 mM KCH3COO, 2 mM Mg(CH3COO)2, 1 mM EGTA, 10% glycerol, pH = 7.6), supplemented with 1 mg/mL BSA, 0.2 mg/mL K-casein, and oxygen scavenging system composed of protocatechuic acid(PCA)/protocatechuate-3,4-dioxygenase(PCD)/Trolox) was used.

For microtubule binding assays, TIRF chambers were first coated with 0.5 mg/ml PLL-PEG-biotin (Surface Solutions Group) followed by with 0.5mg/ml streptavidin (ThermoFisher Scientific). Diluted microtubules were then flowed into the chambers. After addition of each component, chambers were incubated for 5 minutes at room temperature. Once it was confirmed that the microtubules were adhered and polymerized sufficiently, the unbound microtubules were removed by washing the chambers with SRP90 assay buffer. To analyze kinesin-1 recruitment onto the microtubules, the purified motor protein was diluted to 40 nM in the assay buffer with 2 mM 50-adenylylimidodiphosphate (AMP-PNP) (Roche Diagnostics GmbH). Kinesin-1 was combined with each UNC-83 construct separately at a 1:5 molar ratio and incubated on ice for 30 minutes. Following the incubation, the protein mixes were diluted with the assay buffer to get 40 nM kinesin-1 and 200 nM UNC-83 construct in the final solution. As controls, each purified UNC-83 construct was diluted to 200 nM in the assay buffer with 2 mM of AMP-PNP. The solutions were then flowed into the TIRF chambers, one at a time, and incubated for 10 minutes at room temperature to let the proteins interact with AMP-PNP and microtubules. Sequential images of 405 nm, 488 nm and 561 nm channels were acquired. Kinesin-1 average fluorescence fold intensity on microtubules were manually measured using ImageJ (Fiji).

For gliding assays, kinesin-1 was diluted in assay buffer to 200 nM and mixed separately with each UNC-83 construct separately at a 1:5 molar ratio and incubated on ice for 30 minutes. Following the incubation, the protein mixes were further diluted with the assay buffer to obtain final concentrations of 200 nM kinesin-1 and 1 µM UNC-83 construct. Controls consisted of UNC-83 constructs alone at 1 µM. The solutions were incubated for 5-10 minutes at room temperature to allow protein adhesion to the slide, after which diluted microtubules were introduced. Following a 5-minute incubation, unbound proteins and microtubules were washed off using assay buffer containing 2 mM ATP. Microtubules were imaged for 1 minute using 405 nm laser at 10% power and images were taken at 1 frame per second. Single images of 561 nm and 488 nm channels were captured using 15% laser power. Kymographs were generated using ImageJ (Fiji), and microtubule velocities were determined using ImageJ Velocity Measurement Tool plug-in^50^.

TIRF imaging was performed on a Nikon TE microscope equipped with a1.49 N.A. PlanApo 100x objective, a TIRF illuminator (LUN4), and an Andor iXon EMCCD camera. Images were acquired using NIS Elements software (Nikon).

## QUANTIFICATION AND STATISTICAL ANALYSIS

All statistical analyses were performed using GraphPad Prism version 10. Data from hyp7 and P-cell nuclear migration assays were visualized as scatter plots, with means and 95% CI represented as error bars. Sample sizes and specific statistical tests are detailed in the figure legends. Unpaired student *t*-tests were conducted for the indicated comparisons.

### Supplemental Figure Legends

**Figure S1:**
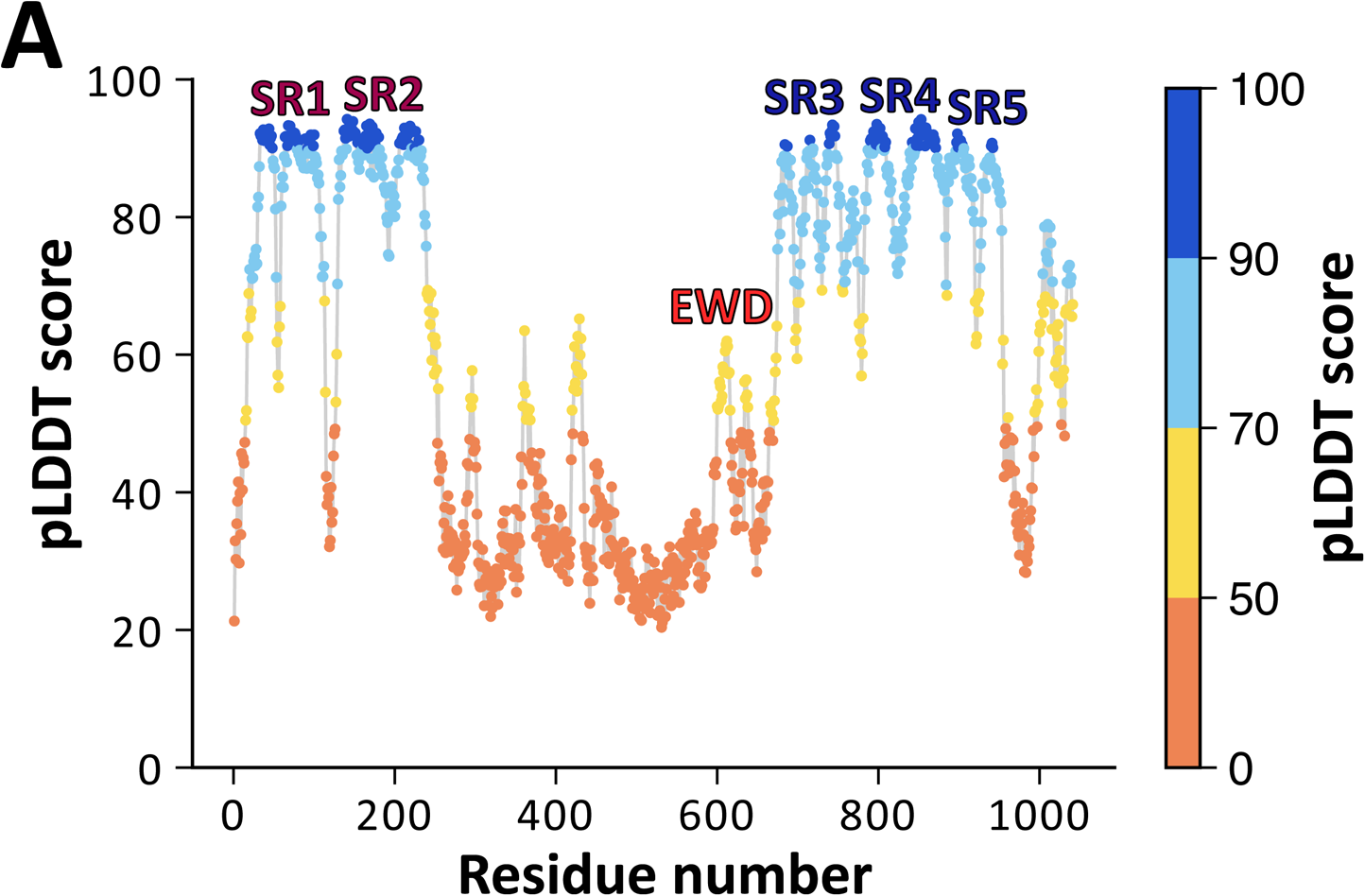
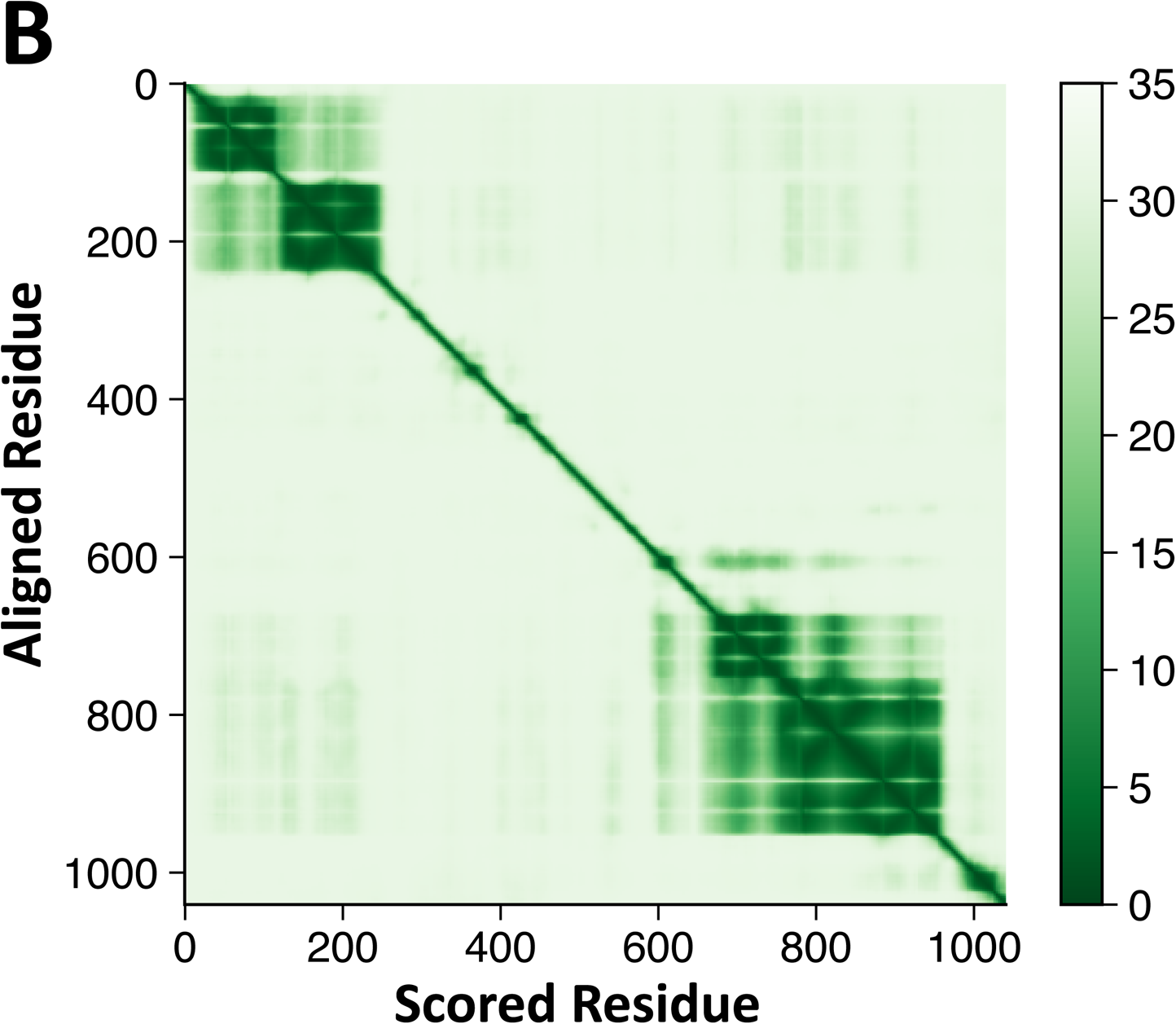
Alphafold3-predicted structure of UNC-83a shows high confidence for spectrin-like repeats, related to Figure 1. A) Plot of the predicted local distance difference test (pLDDT) scores per residue; high confidence in dark blue, low confidence in orange/yellow. B) Plot of the predicted aligned error (PAE) per residue; higher error in light green.

**Figure S2:**
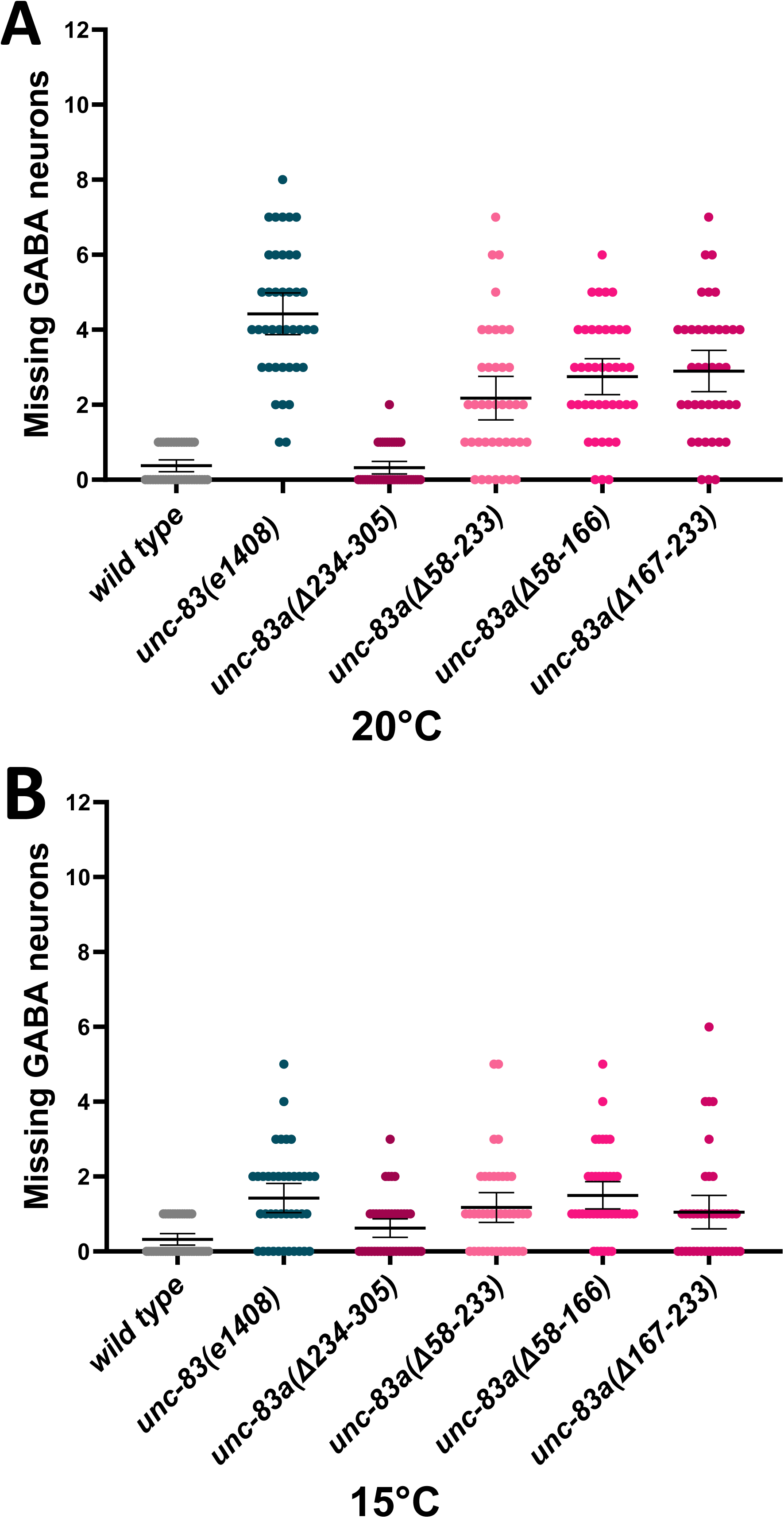
Deleting spectrin-like repeats in the UNC-83a-specific N-terminal domain causes temperature-sensitive P-cell nuclear migration defects, related to Figure 4. Quantification of larval P-cell nuclear migration in *wild type*, *unc-83(e1408)* null, and *unc-83a* N-terminal domain deletion mutant animals (A) at 20°C and (B) at 15°C. Each point represents the number of missing GABA neurons per animal and n = 40 for each strain.

**Figure S3:**
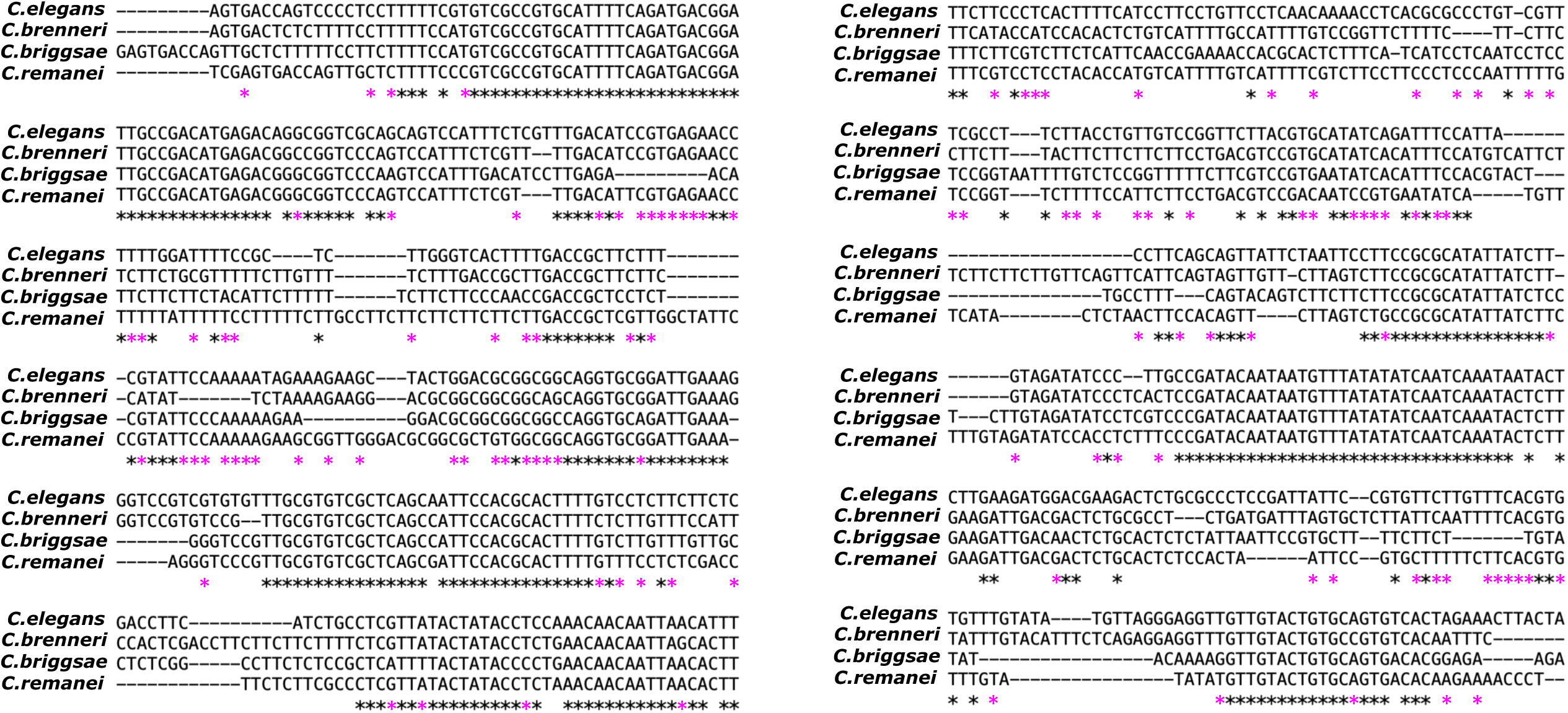
The *unc-83* enhancer region shows high conservation across *Caenorhabditis* species, related to Figure 5. Multiple sequence alignment comparing the long intron region of *C. elegans unc-83* with corresponding sequences from *C. brenneri*, *C. remanei* and *C. briggsae*. Black asterisks indicate nucleotides conserved across all four species while magenta asterisks indicate that the *C. elegans* nucleotides conserved in two other species.

**Movie S1: *C. elegans* kinesin-1 heterotetramer demonstrates microtubule gliding activity, related to Figure 6**. Representative time-lapse showing microtubules (blue) gliding across a glass surface. Scale bar (= 5 µm) and time stamp are indicated in the movie.

**Movie S2: The *C. elegans* kinesin-1 heterotetramer preincubated with UNC-83c increases the gliding velocity of microtubules, related to Figure 6**. Representative time-lapse showing microtubules (blue) gliding across a glass surface when kinesin-1 heterotetramer is preincubated with UNC-83c. Scale bar (= 5 µm) and time stamp are indicated in the movie.

**Movie S3: The *C. elegans* kinesin-1 heterotetramer preincubated with UNC-83a increases the gliding velocity of microtubules, related to Figure 6**. Representative time-lapse showing microtubules (blue) gliding across a glass surface when kinesin-1 heterotetramer is preincubated with UNC-83a. Scale bar (= 5 µm) and time stamp are indicated in the movie.

**Movie S4: UNC-83a N-terminus inhibits *C. elegans* kinesin-1 heterotetramer motor activity, related to Figure 6**. Representative time-lapse showing reduced microtubule (blue) gliding across a glass surface when kinesin-1 heterotetramer is preincubated with UNC-83a^N-term^. Scale bar (= 5 µm) and time stamp are indicated in the movie.

---

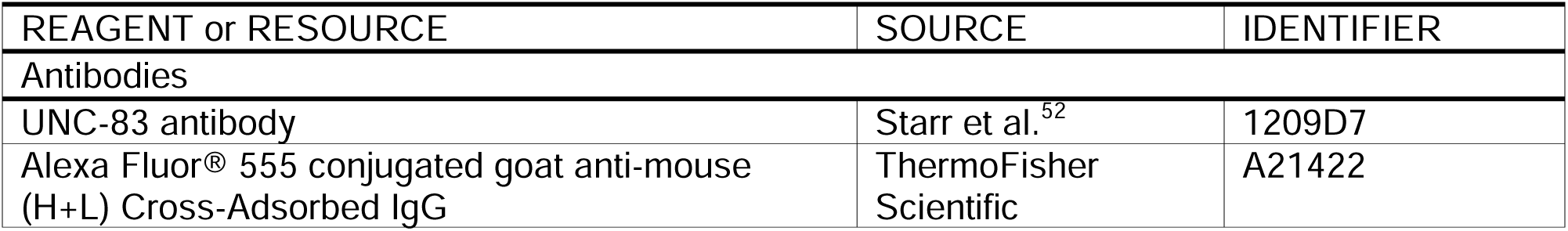
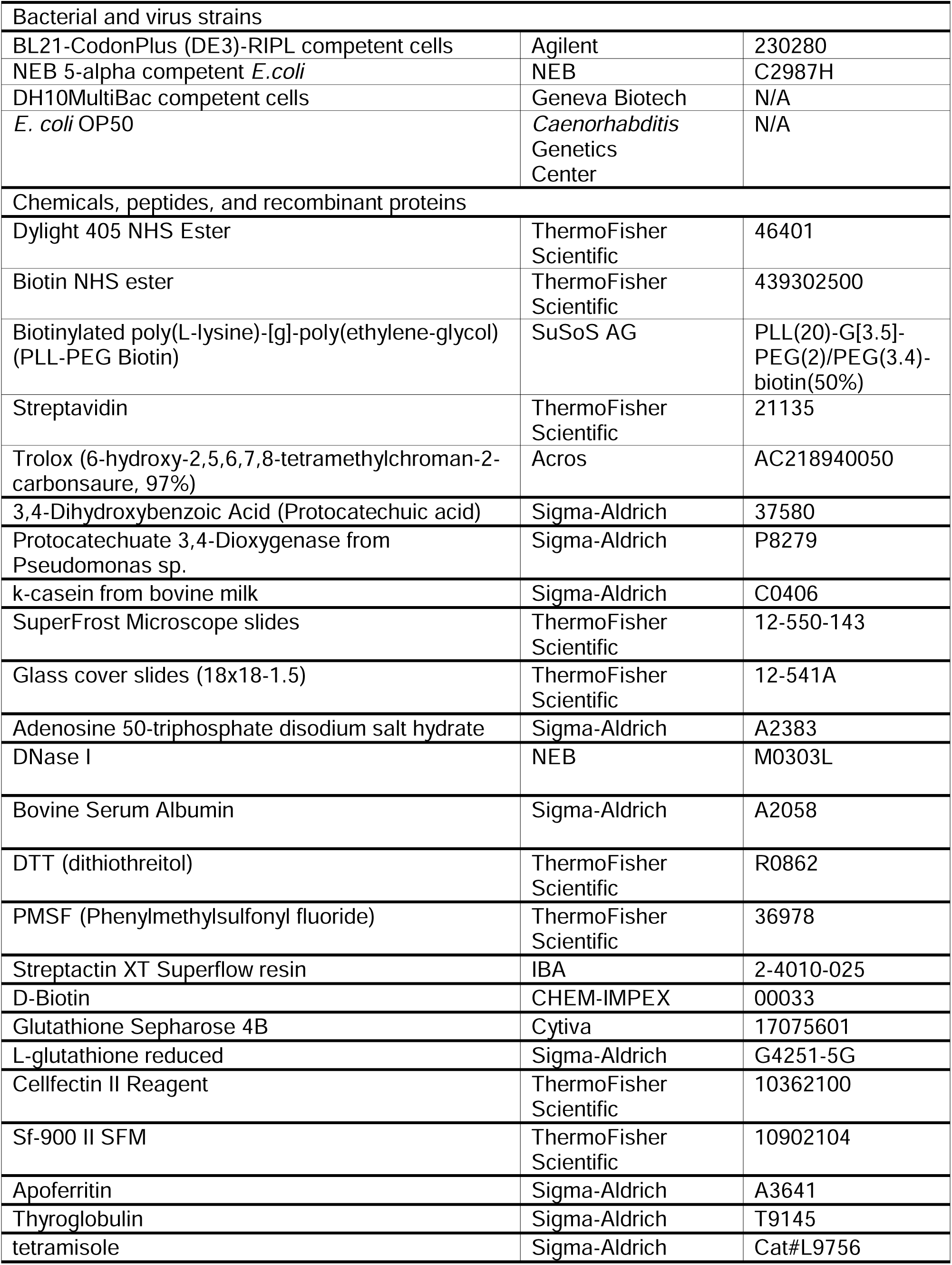
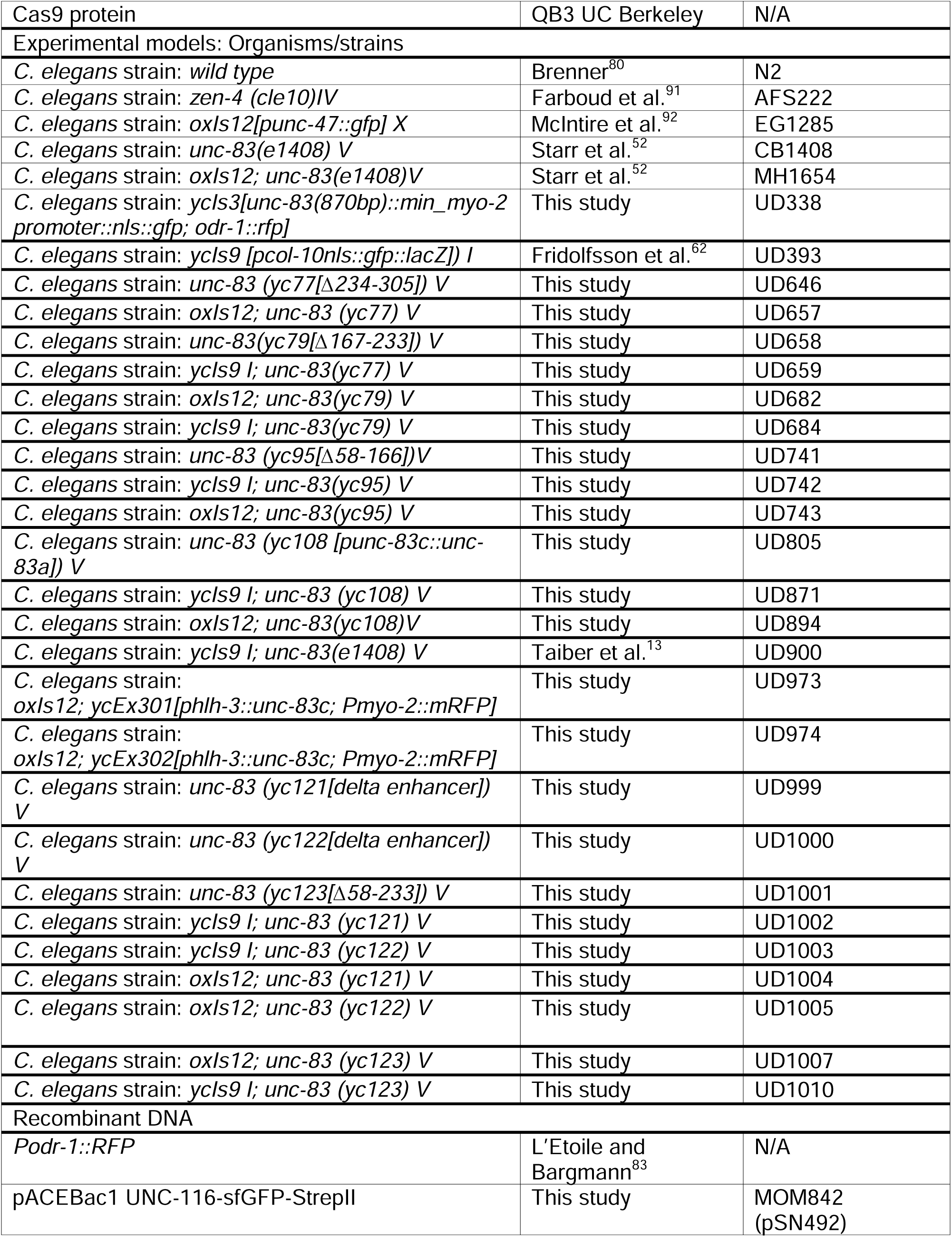
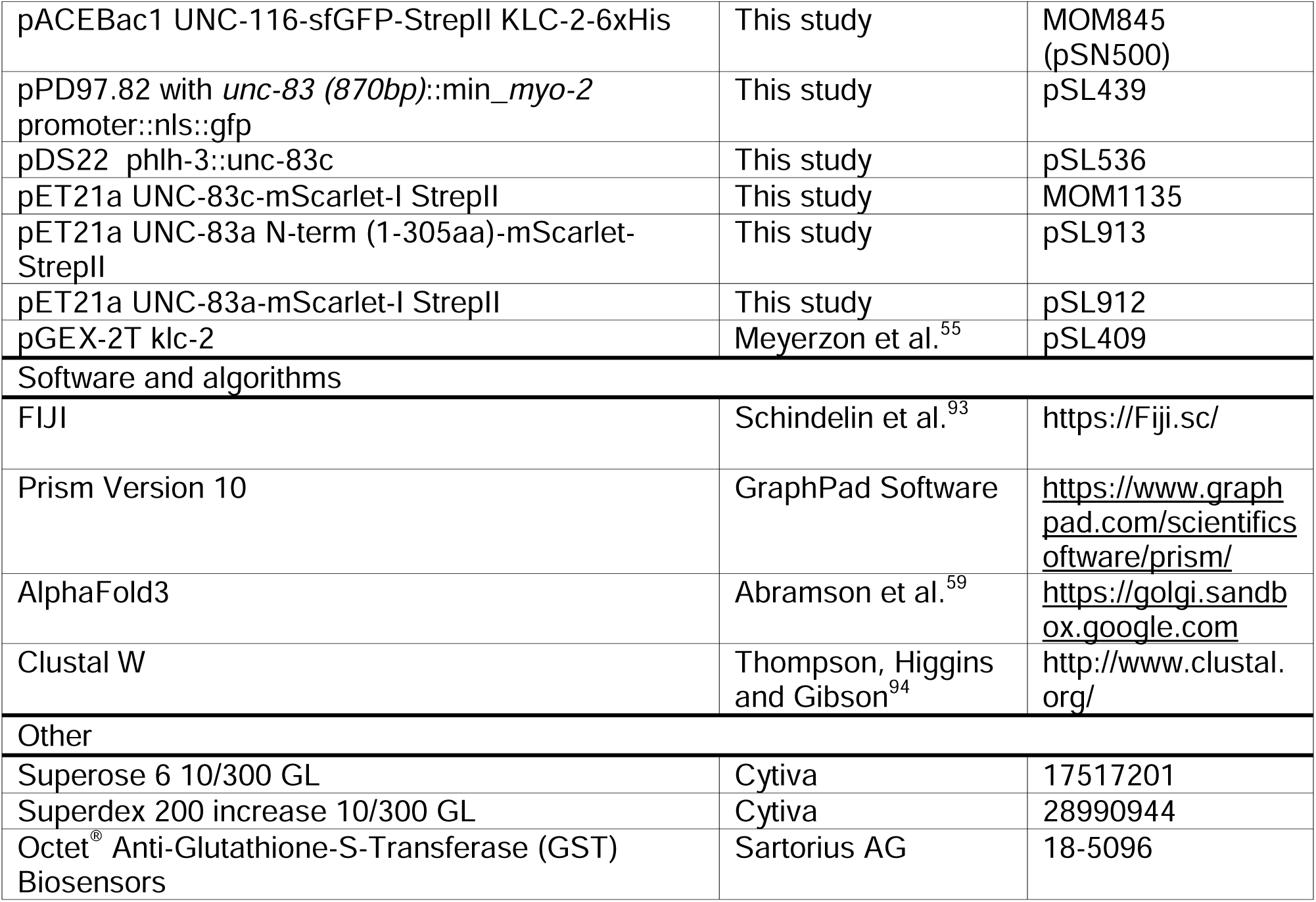
Key Resources Table.

**Table S1.**
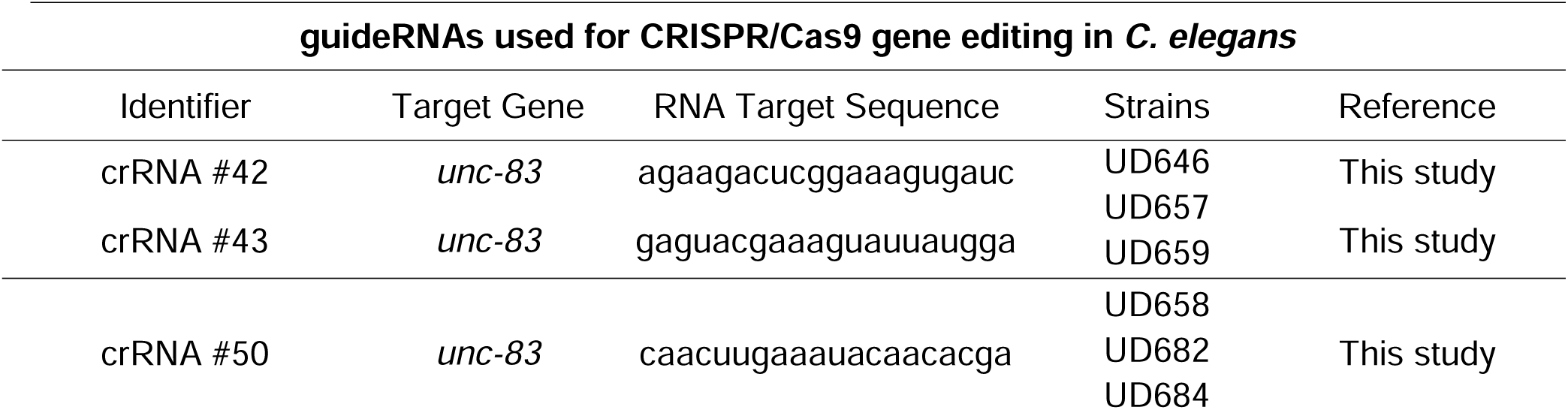
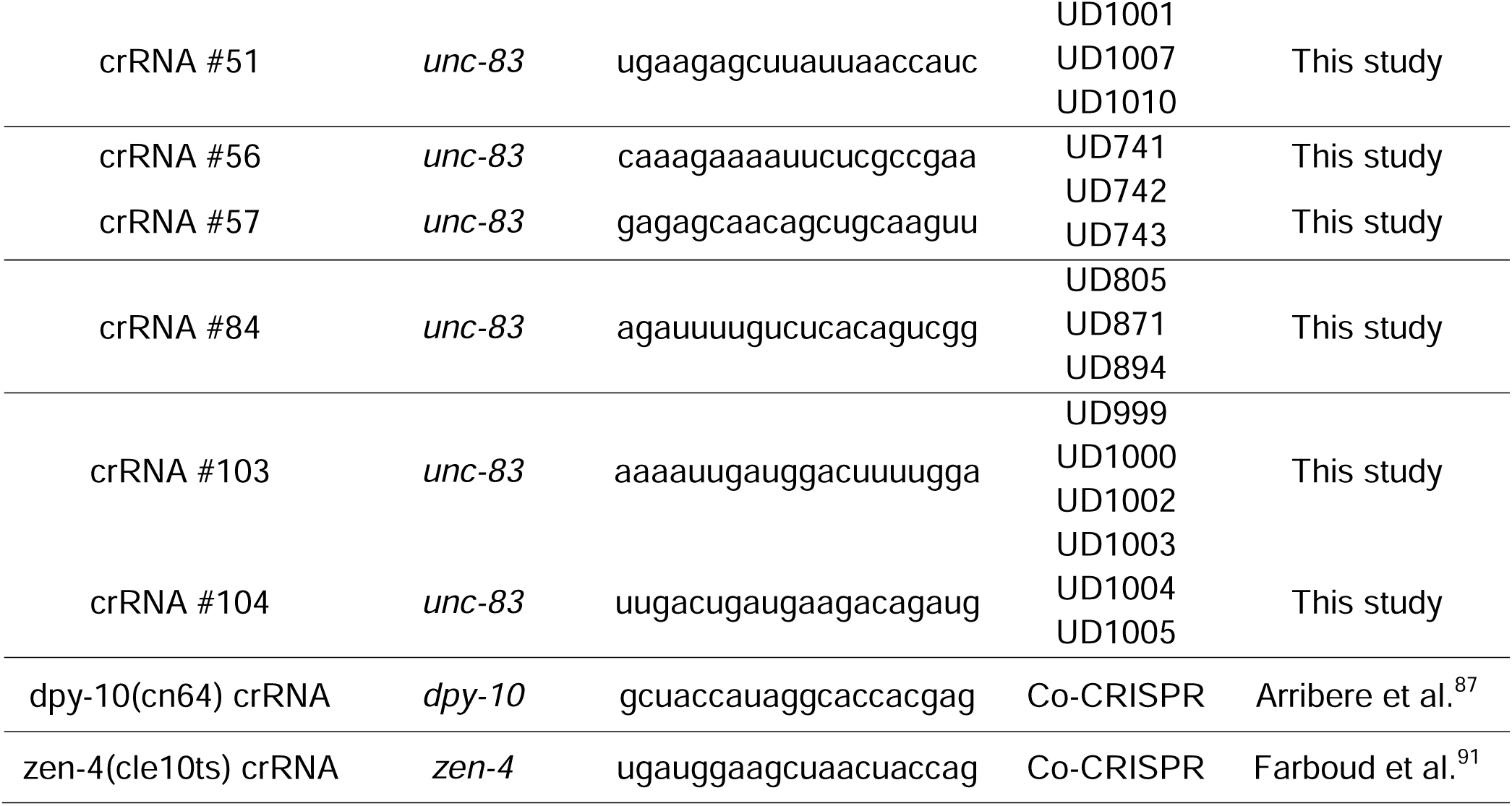
guideRNAs used in this study. Related to STAR Methods.

**Table S2.**
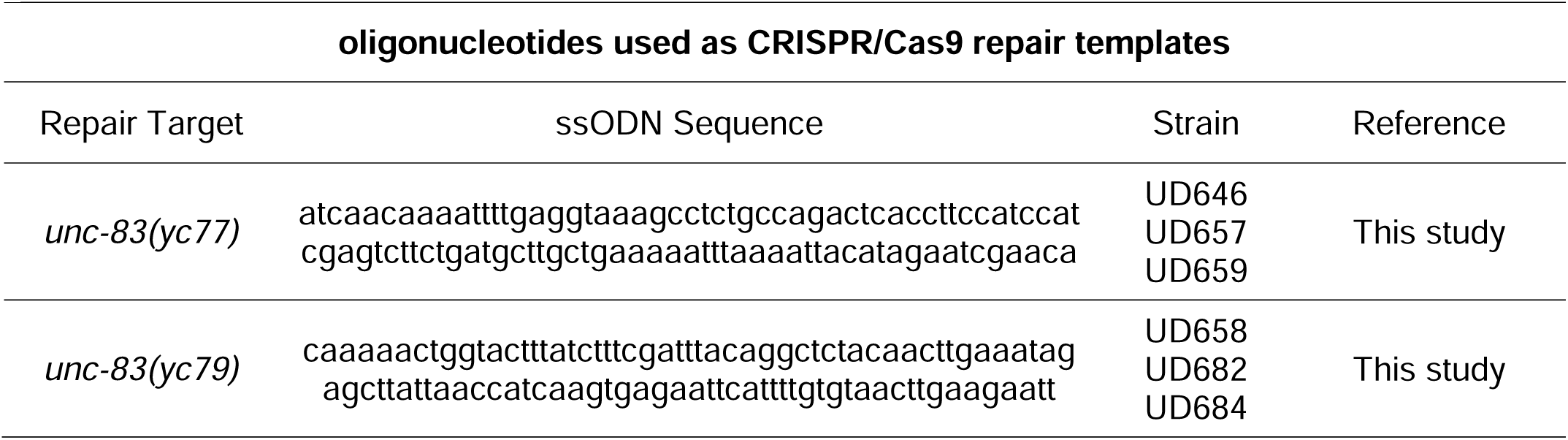
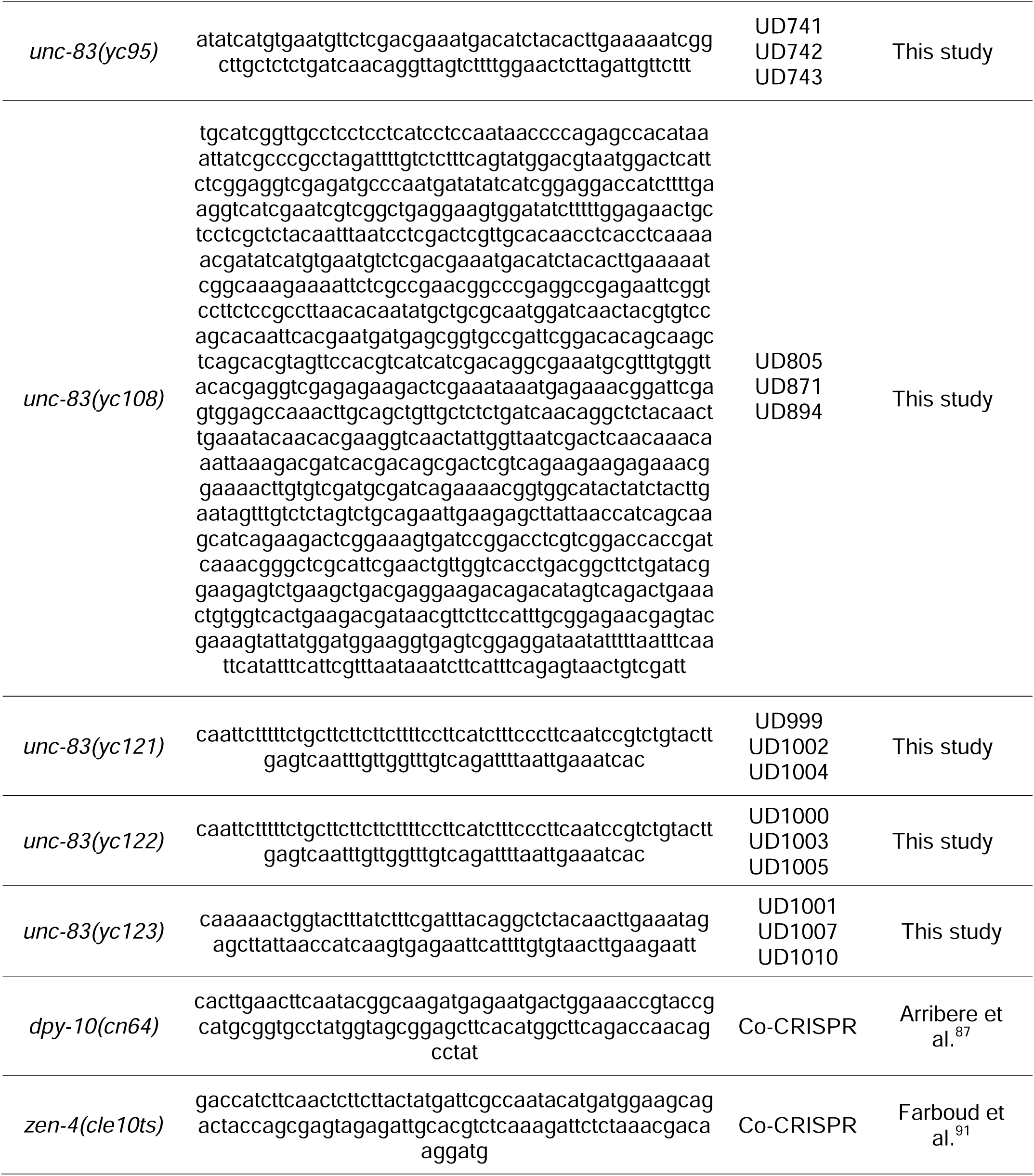
CRISPR/Cas-9 repair templates used in this study. Related to STAR Methods.

